# Dynamic targeting enables domain-general inhibitory control over action and thought by the prefrontal cortex

**DOI:** 10.1101/2020.10.22.350520

**Authors:** Dace Apšvalka, Catarina S. Ferreira, Taylor W. Schmitz, James B. Rowe, Michael C. Anderson

## Abstract

Successful self-control requires the ability to stop unwanted actions or thoughts. Stopping is regarded as a central function of inhibitory control, a mechanism enabling the suppression of diverse mental content, and strongly associated with the prefrontal cortex. A domain-general inhibitory control capacity, however, would require the region or regions implementing it to dynamically shift top-down inhibitory connectivity to diverse target regions in the brain. Here we show that stopping unwanted thoughts and stopping unwanted actions engage common regions in the right anterior dorsolateral and right ventrolateral prefrontal cortex, and that both areas exhibit this dynamic targeting capacity. Within each region, pattern classifiers trained to distinguish stopping actions from making actions also could identify when people were suppressing their thoughts (and vice versa) and could predict which people successfully forgot thoughts after inhibition. Effective connectivity analysis revealed that both regions contributed to action and thought stopping, by dynamically shifting inhibitory connectivity to motor area M1 or to the hippocampus, depending on the goal, suppressing task-specific activity in those regions. These findings support the existence of a domain-general inhibitory control mechanism that contributes to self-control and establish dynamic inhibitory targeting as a key mechanism enabling these abilities.

## Introduction

Well-being during difficult times requires the ability to stop unwelcome thoughts. This vital ability may be grounded in inhibitory control mechanisms that also stop physical actions (Anderson & Hanslmayr, 2014; Anderson et al., 2004; Castiglione et al., 2019; Depue et al., 2016; Depue et al., 2007). According to this hypothesis, the right lateral prefrontal cortex (rLPFC) supports self-control, allowing people to regulate their thoughts and behaviours when fears, ruminations, or impulsive actions might otherwise hold sway (Anderson & Hulbert, 2021; Benoit et al., 2016; Schmitz et al., 2017). This proposal rests on the concept of inhibitory control, a putative domain-general control mechanism that has attracted much interest in psychology and neuroscience over the last two decades (Anderson et al., 2016; Aron et al., 2004, 2014; Banich & Depue, 2015; Bari & Robbins, 2013; Boucher et al., 2007; Diamond, 2013; Ersche et al., 2012; Eysenck et al., 2007; Joormann & Tanovic, 2015; Lipszyc & Schachar, 2010). Despite widespread and enduring interest, central evidence for the neural basis of domain-general inhibitory control is missing: no study has shown a control region that dynamically shifts its connectivity to suppress local processing in diverse cortical areas depending on the stopping goal – a fundamental capability of this putative mechanism. Inhibiting actions and memories, for example, requires that an inhibitory control region target disparate specialised brain areas to suppress motoric or mnemonic processing, respectively. We term this predicted capability dynamic targeting. Here, we tested the existence of dynamic targeting by asking participants to stop unwanted actions or thoughts. Using functional magnetic resonance imaging (fMRI) and pattern classification, we identified prefrontal regions that contribute to successful stopping in both domains. Critically, we then tested whether people’s intentions to stop actions or thoughts were reflected in altered effective connectivity between the domain-general inhibition regions in prefrontal cortex with memory or motor-cortical areas. By tracking the dynamic targeting of inhibitory control in the brain, we provide a window into humans’ capacity for self-control over their thoughts and behaviours (Nigg, 2017).

Our analysis builds on evidence that two regions of the rLPFC may contribute to stopping both actions and thoughts: the right ventrolateral prefrontal cortex (rVLPFC) and the right dorsolateral prefrontal cortex (rDLPFC). For example, stopping motor actions activates rVLPFC (especially in BA44/45, pars opercularis), rDLPFC, and anterior insula (Aron et al., 2004; Guo et al., 2018; Jahanshahi et al., 2015; Levy & Wagner, 2011; Rae et al., 2014; Zhang et al., 2017). Disrupting rVLPFC impairs motor inhibition, whether via lesions (Aron et al., 2003), transcranial magnetic stimulation (Chambers et al., 2006), intracranial simulation in humans (Wessel et al., 2013) or monkeys (Sasaki et al., 1989). RVLPFC thus could promote top-down inhibitory control over actions, and possibly inhibitory control more broadly (Aron, 2007; Aron et al., 2004; Castiglione et al., 2019). Within-subjects comparisons have identified shared activations in rDLPFC (BA 9/46) that could support a domain-general mechanism that stops both actions and thoughts (Depue et al., 2016).

If these rLPFC regions support domain-general inhibitory control, the question arises as to how inhibition is directed at actions or thoughts. To address this issue, we tested whether any regions within the rLPFC had the dynamic targeting capacity needed to support domain-general inhibitory control. Dynamic targeting requires that a candidate inhibitory control system exhibit five core attributes (see Figure 1). First, stopping in diverse domains should engage the proposed source of control, with activation patterns within this region transcending the specific demands of each stopping type. As a consequence, activation patterns during any one form of stopping should contain information shared with inhibition in other domains. Second, the engagement of the proposed prefrontal source should track indices of inhibitory control in diverse domains, demonstrating its behavioural relevance. Third, stopping-related activity in the prefrontal sources should co-occur with interrupted functioning in domain-specific target sites representing thoughts or actions. Fourth, the prefrontal source should exert top-down inhibitory coupling with these target sites, providing the causal basis of their targeted suppression. Finally, dynamic targeting requires that inhibitory coupling between prefrontal source and domain-specific target regions be selective to current goals.

**Figure 1.**
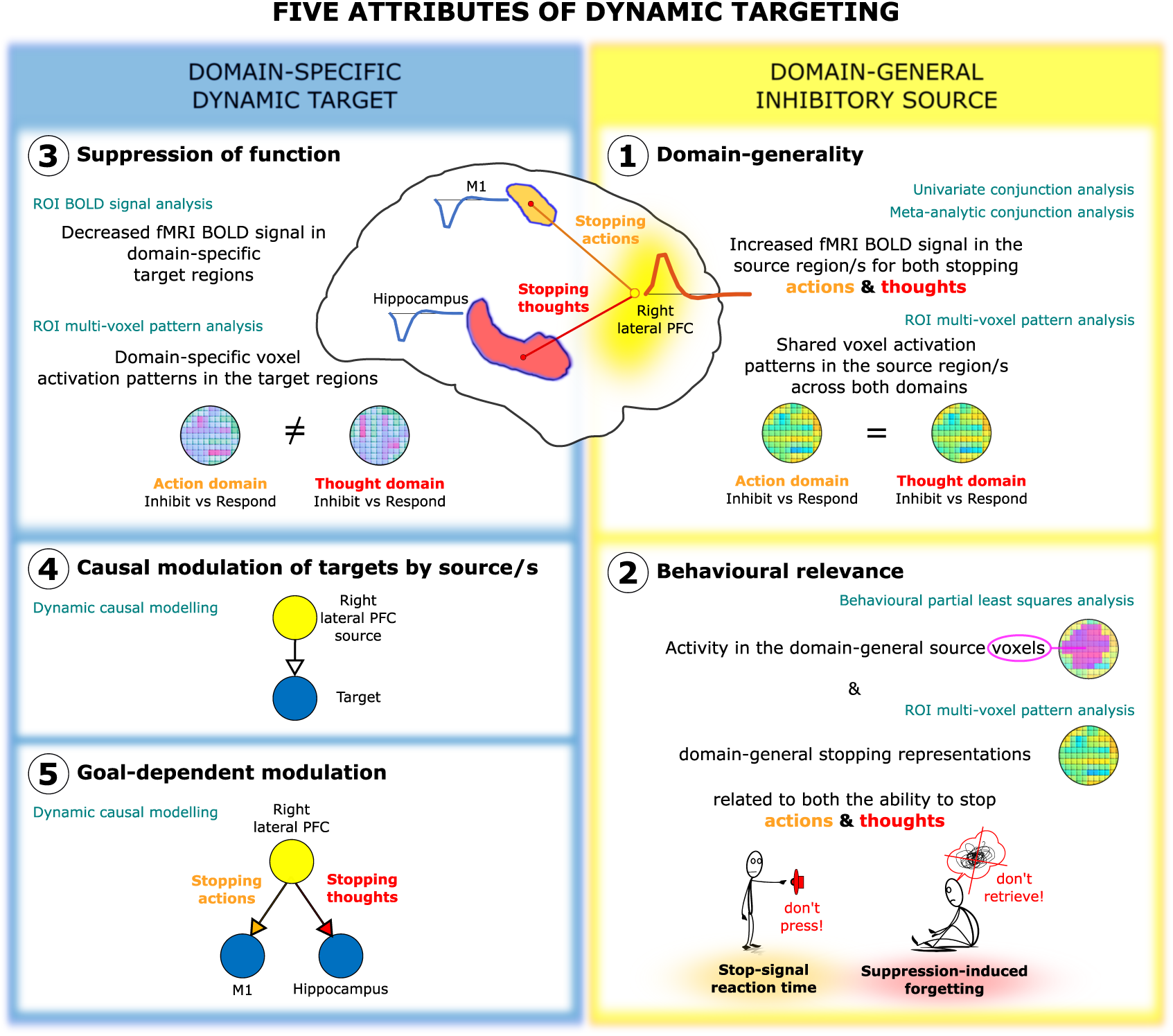
Five attributes of dynamic targeting. Schematic of the five attributes of domain-general inhibitory control by dynamic targeting and methods employed (teal colour) to test the attributes. Attributes 1-2 relate to the existence of domain-general inhibitory sources. The predicted location of such sources was in the right lateral PFC. We present the two attributes on the right side to match the visualised location of the expected sources. To test the domain-generality of inhibitory sources (attribute 1), we performed univariate and meta-analytic conjunction analysis of the No-Think > Think and Stop > Go contrasts, and cross-task multi-voxel pattern analysis (MVPA). To test the behavioural relevance (attribute 2), we related inhibitory activations within the identified domain-general regions to individual variation in inhibition ability (stop-signal reaction time and suppression-induced forgetting) using behavioural partial-least squares and MVPA. Attributes 3-5 relate to the existence of domain-specific target sites that are dynamically modulated by the domain-general sources. Our a priori assumption was that suppressing actions and thoughts would target M1 and hippocampus, respectively. To test the suppression of function within the target sites (attribute 3) we performed a region of interest (ROI) analysis expecting down-regulation within the target sites, and cross-task MPVA expecting distinct activity patterns across the two task domains. To test whether the prefrontal domain-general sources exert top-down modulation of the target sites (attribute 4) dynamically targeting M1 or the hippocampus depending on the process being stopped (attribute 5), we performed dynamic causal modelling.

These attributes of dynamic targeting remain unproven, despite the fundamental importance of inhibitory control. Research on response inhibition and thought suppression instead has focused on how the prefrontal cortex contributes to stopping within each domain (Anderson et al., 2016; Jana et al., 2020; Schall et al., 2017; Wiecki & Frank, 2013). For example, research on thought suppression has revealed top-down inhibitory coupling from the rDLPFC to the hippocampus, and to several cortical regions representing specific mnemonic content (Benoit & Anderson, 2012; Benoit et al., 2015; Gagnepain et al., 2014; Gagnepain et al., 2017; Mary et al., 2020; Schmitz et al., 2017). Moreover, suppressing thoughts down-regulates hippocampal activity, with the down-regulation linked to hippocampal GABA and forgetting of the suppressed content (Schmitz et al., 2017). Top-down modulation of actions by rVLPFC suggests that premotor and primary motor cortex are target sites (Aron & Poldrack, 2006; Rae et al., 2015; Zandbelt et al., 2013). Action stopping engages local intracortical inhibition within M1 to achieve stopping (Coxon et al., 2006; Sohn et al., 2002; Stinear et al., 2009; van den Wildenberg et al., 2010), with a person’s stopping efficacy related to local GABAergic inhibition (He et al., 2019). However, studies of thought suppression and action stopping posit that control originates from different prefrontal regions (rDLPFC vs rVLPFC), possibly reflecting domain-specific inhibitory control mechanisms. A candidate source of domain-general inhibitory control must stop both actions and thoughts and exhibit the attributes of dynamic targeting.

Although dynamic inhibitory targeting has not been tested, some large-scale networks flexibly shift their coupling with diverse brain regions that support task performance. Diverse tasks engage a fronto-parietal network (Cole et al., 2013; Cole & Schneider, 2007; Duncan, 2010; Fox et al., 2005), which exhibits greater cross-task variability in coupling with other regions than other networks (Cocuzza et al., 2020; Cole et al., 2013). Variable connectivity may index this network’s ability to reconfigure flexibly and coordinate multiple task elements in the interests of cognitive control (Cole et al., 2013). A cingulo-opercular network, including aspects of rDLPFC and rVLPFC, also is tied to cognitive control, including conflict and attentional processing (Botvinick, 2007; Cole et al., 2009; Crittenden et al., 2016; Dosenbach et al., 2006; Petersen & Posner, 2012; Seeley et al., 2007; Yeo et al., 2015), with the prefrontal components exhibiting high connectivity variability over differing tasks (Cocuzza et al., 2020). However, previous analyses of these networks do not address dynamic inhibitory targeting: Dynamic targeting requires not merely that the prefrontal cortex exhibits connectivity to multiple regions, but that the connectivity includes a top-down component that suppresses target regions.

We sought to test the presence of dynamic targeting through the properties of prefrontal, motor and hippocampal networks (see Figure 1 for an overview of our approach). We combined, within one fMRI session, a cognitive manipulation to suppress unwanted thoughts, the Think/No-Think paradigm (Anderson & Green, 2001; Anderson & Hulbert, 2021), with motor action stopping in a stop-signal task (Logan & Cowan, 1984; Verbruggen et al., 2019). This design provided the opportunity to identify colocalized activations of domain-general inhibitory control in prefrontal sources and observe their changes in effective connectivity with motor cortical and hippocampal targets. For the thought suppression task, prior to scanning, participants learned word pairs, each composed of a reminder and a paired thought (Figure 2). During thought stopping scanning blocks, on each trial, participants viewed one of these reminders. For each reminder, we cued participants either to retrieve its associated thought (Think trials) or instead to suppress its retrieval, stopping the thought from coming to mind (No-Think trials). For the action stopping task, prior to scanning, participants were trained to press one of two buttons in response to differently coloured circles (Schmitz et al., 2017). During the action stopping scanning blocks, participants engaged in a speeded motor response task that, on a minority of trials, required them to stop their key-press following an auditory stop signal. Action and thought stopping blocks alternated, to enable quantification of domain-general and domain-specific activity and connectivity.

**Figure 2.**
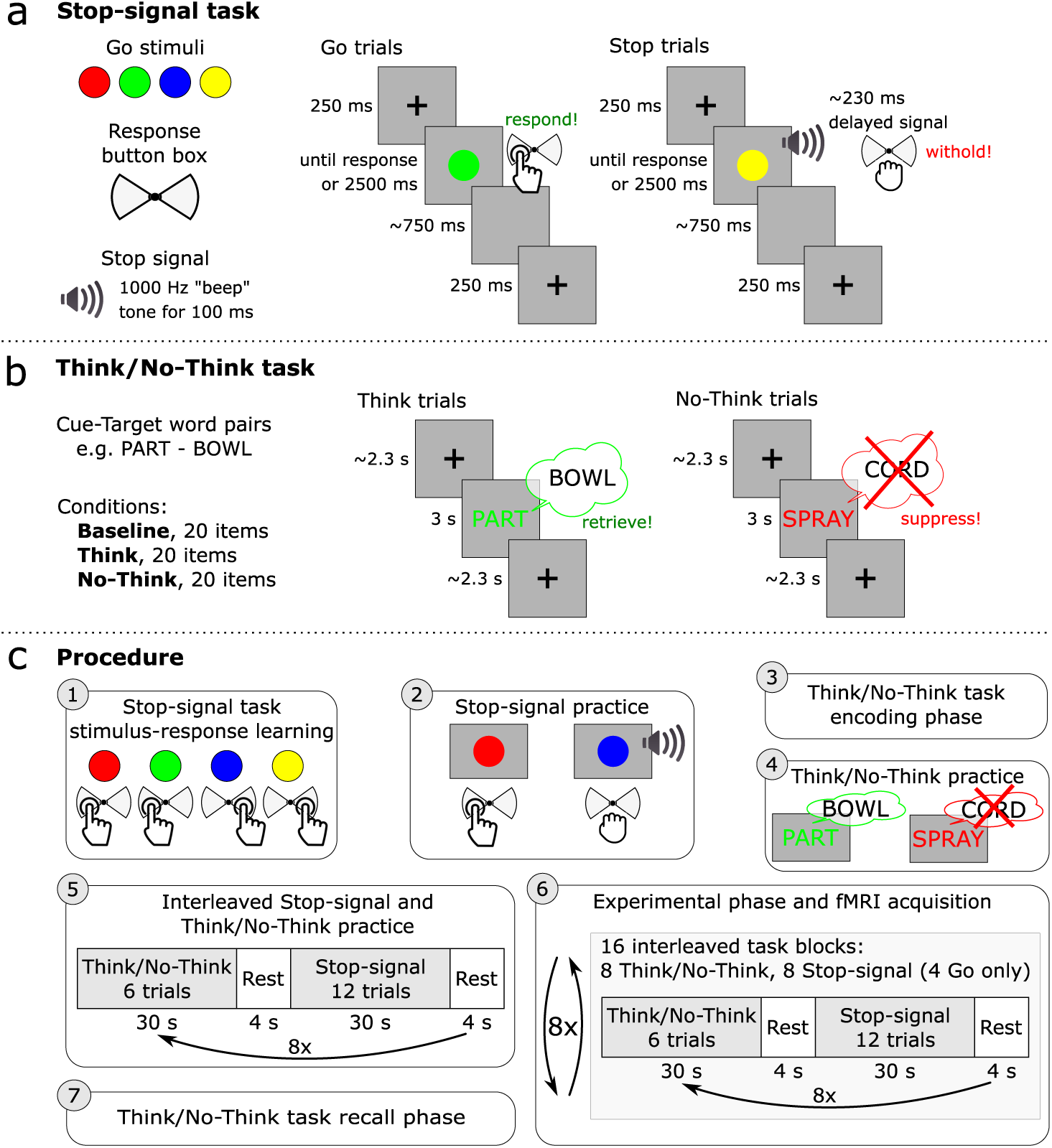
Schematic of the experimental paradigm and procedure. **(a)** In the Stop-signal task, the Go stimuli were red, green, blue, and yellow coloured circles. On Go trials, participants responded by pressing one of the two buttons on a button box according to learned stimulus-response associations. On Stop trials, shortly after the Go stimulus, an auditory “beep” tone would signal participants to withhold the button press. The stop-signal delay varied dynamically in 50 ms steps to achieve approximately a 50% success-to-stop rate for each participant. **(b)** In the Think/No-Think task, participants learned 78 cue-target word pair associations. Sixty of the word pairs were then divided into three lists composed of 20 items each and allocated to the three experimental conditions: Think, No-Think, and Baseline. During Think trials, a cue word appeared in green, and participants had 3 s to retrieve and think of the associated target word. On No-Think trials, a cue word appeared in red and participants were asked to suppress the retrieval of the associated target word and push it out of awareness if it popped into their mind. **(c)** The procedure consisted of 7 steps: 1) stimulus-response learning for the Stop-signal task: 2) Stop-signal task practice; 3) encoding phase of the Think/No-Think task; 4) Think/No-Think practice; 5) practice of interleaved Stop-signal and Think/No-Think tasks; 6) the main experimental phase during fMRI acquisition where participants performed interleaved 30 s blocks of Stop-signal and Think/No-Think tasks; 7) recall phase of the Think/No-Think task.

The dynamic targeting hypothesis predicts that stopping actions and thoughts call upon a common inhibition mechanism. For thought suppression, we predicted that the reminder would activate the associated thought, triggering inhibitory control to suppress hippocampal retrieval (Anderson et al., 2004; Levy & Anderson, 2012). We predicted that this disruption would hinder later retrieval of the thought, causing suppression-induced forgetting. To verify this, we tested all pairs (both Think and No-Think pairs) after scanning, including a group of pairs that had been learned, but that were omitted during the Think/No-Think task, to estimate baseline memory performance (Baseline pairs). Suppression-induced forgetting occurs when final recall of No-Think items is lower than Baseline items (Anderson & Green, 2001). For action stopping, we proposed that the Go stimulus would rapidly initiate action preparation, with the presentation of the stop signal triggering inhibitory control to suppress motor processes in M1 (Logan & Cowan, 1984; Verbruggen et al., 2019). If the capacities to stop actions and thoughts are related, more efficient action stopping, as measured by stop-signal reaction time, should correlate with greater suppression-induced forgetting.

Our primary goal was to determine whether any prefrontal source region meets the five core attributes for dynamic targeting of inhibitory control. To test this, we first identified candidate regions that could serve as sources of control. We isolated prefrontal regions that were more active during action and thought stopping, compared to their respective control conditions (e.g. “Go” trials, wherein participants made the cued action; or Think trials, wherein they retrieved the cued thought) and then performed a within-subjects conjunction analysis on these activations. We performed a parallel conjunction analysis on independent data from two quantitative meta-analyses of fMRI studies that used the Stop-signal or the Think/No-Think tasks, to confirm the generality of the regions identified. We next tested whether activation patterns within these potential source regions transcended the particular stopping domains. We used multi-voxel activation patterns to train a classifier to discriminate stopping from going in one modality (e.g., action stopping), to test whether it could identify stopping in the other modality (e.g. thought suppression). Finally, to examine behavioural relevance, we related inhibitory activations within these meta-analytic conjunction areas to individual variation in inhibition ability (e.g., suppression-induced forgetting and stop-signal reaction time) using behavioural partial least squares and multi-voxel pattern analysis. Any regions surviving these constraints was considered a strong candidate for a hub of inhibitory control. We hypothesized that these analyses would identify the right anterior DLPFC (Anderson & Hulbert, 2021; Benoit & Anderson, 2012; Depue et al., 2016; Guo et al., 2018), and right VLPFC (Aron et al., 2004; Levy & Wagner, 2011).

To verify that inhibitory control targets goal-relevant brain regions, we next confirmed that *a priori* target sites are suppressed in a goal-specific manner. Specifically, stopping retrieval should down-regulate hippocampal activity (Anderson et al., 2004; Benoit & Anderson, 2012; Depue et al., 2007; Gagnepain et al., 2014; Gagnepain et al., 2017; Levy & Anderson, 2012; Mary et al., 2020), more than does action stopping. In contrast, stopping actions should inhibit motor cortex more than does thought stopping (Schmitz et al., 2017). To determine whether these differences in modulation arise from inhibitory targeting by our putative domain-general prefrontal control regions, we used dynamic causal modelling (Friston et al., 2003). If both DLPFC and VLPFC are involved, as prior work suggests, we sought to evaluate whether one or both of these regions are critical sources of inhibitory control.

## Results

### The ability to inhibit unwanted thoughts is related to action stopping efficiency

We first tested whether action stopping efficiency was associated to successful thought suppression. To quantify action stopping efficiency, we computed stop-signal reaction times (SSRTs) using the consensus standard integration method (Verbruggen et al., 2019). We confirmed that the probability of responding to Stop trials (M = 0.49, SD = 0.07; ranging from 0.36 to 0.69) fell within the recommended range for reliable estimation of SSRTs (Verbruggen et al., 2019), and that the probability of Go omissions (M = 0.002, SD = 0.01) and choice errors on Go trials (M = 0.04, SD = 0.02) were low. We next verified that the correct Go RT (M = 600.91 ms, SD = 54.63 ms) exceeded the failed Stop RT (M = 556.92 ms, SD = 56.77) in all but one participant (9 ms difference between the failed Stop RT and correct Go RT; including this participant makes little difference to any analysis, so they were not excluded). Given that the integration method requirements were met, the average SSRT, our measure of interest, was 348.34 ms (SD = 51.25 ms), with an average SSD of 230 ms (SD = 35.68 ms).

We next verified that our Think/No-Think task had induced forgetting of suppressed items. We compared final recall of No-Think items to that of Baseline items that had neither been suppressed nor retrieved (see Methods). Consistent with a previous analysis of these data (Schmitz et al., 2017) and with prior findings (Anderson & Green, 2001; Anderson & Huddleston, 2012; Anderson et al., 2004; Levy & Anderson, 2012), suppressing retrieval impaired No-Think recall (M = 72%, SD = 9%) relative to Baseline recall (M = 77%, SD = 9%), yielding a suppression-induced forgetting (SIF) effect (Baseline – No-Think = 5%, SD = 9%, one-tailed t_23_ = 2.55, p = 0.009, d = 0.521). Thus, suppressing retrieval yielded the predicted inhibitory aftereffects on unwanted thoughts.

To test the relationship between thought suppression and action stopping, we calculated a SIF score for each participant by subtracting No-Think from Baseline recall performance (Baseline – No-think). This metric indexes the efficiency with which each participant could down-regulate later accessibility of suppressed items, an after-effect of suppression believed to be sensitive to inhibitory control (Anderson & Green, 2001). We then correlated the SSRT and SIF scores (excluding one bi-variate outlier; see Methods). Consistent with a potential shared inhibition process, better action stopping efficiency (faster SSRTs) was associated with greater SIF (r_ss_ = −0.492, p = 0.014, see Figure 4a; A detailed report of behavioural results is available in the supplementary analysis notebook).

Although we quantified SSRT with the integration method, this method may, at times, overestimate SSRTs because it does not consider times when participants fail to trigger the stopping process, known as trigger failures (Matzke et al., 2017). Trigger failures may arise, for example, when a participant is inattentive and misses a stop signal. We recomputed SSRTs using a method that estimates trigger failure rate and that corrects SSRTs for these events (Matzke et al., 2017; Matzke et al., 2013). This method yielded shorter SSRTs (M = 278.84 ms, SD = 41.13 ms) than the integration method (M = 348.34 ms), but did not alter the relationship between stopping efficiency and SIF (r = −0.383, p = 0.065), which remained similar to the relationship observed with integration method (r_ss_ = −0.492, p = 0.014). This alternate SSRT measure also did not qualitatively alter brain-behaviour relationships reported throughout. These findings suggest that attentional factors that generate trigger failures are unlikely to explain the relationship between thought and action inhibition.

### Stopping actions and memories engages both right DLPFC and VLPFC

We next isolated brain regions that could provide a source of inhibitory control over action and thought. The whole-brain voxel-wise conjunction analysis of the Stop > Go and the No-Think > Think contrasts revealed that both motor and thought inhibition evoked conjoint activations in the right prefrontal cortex (PFC), specifically, the rDLPFC (middle frontal and superior frontal gyri), rVLPFC (ventral aspects of inferior frontal gyrus, including BA44/45, extending into insula), precentral gyrus, and supplementary motor area (see Table 1a and Figure 3). These findings suggest a role of the right PFC in multiple domains of inhibitory control (Aron et al., 2004; Depue et al., 2016; Garavan et al., 1999), a key attribute necessary to establish dynamic targeting.

**Table 1.** Within-subjects and meta-analysis domain-general inhibition-induced activations (Stop > Go & No-Think > Think)

| Nr. | Hemisphere | Region | ~BA | Network | MNI of the peak |  |  | Volume (mm<sup>3</sup>) |
| --- | --- | --- | --- | --- | --- | --- | --- | --- |
|  |  |  |  |  | x | y | z |  |
| a. Within-subjects, Stop >Go & No-Think >Think |  |  |  |  |  |  |  |  |
| 1 | Right | Inferior frontal gyrus (VLPFC)<br>Insula | 44, 45 | CON, FPN | 45 | 18 | 8 | 5366 |
| 2 | Right | Inferior parietal lobule | 40 | CON, FPN, PMM | 63 | -42 | 41 | 3611 |
| 3 | Right | Supplementary motor area | 6, 8 | CON, FPN, LAN | 15 | 18 | 64 | 2498 |
| 4 | Right | Middle frontal gyrus (DLPFC)<br>Superior frontal gyrus (DLPFC) | 9, 10, 46 | CON | 33 | 42 | 23 | 1654 |
| 5 | Right | Precentral gyrus | 6 | CON, FPN, LAN | 42 | 3 | 41 | 945 |
| 6 | Left | Inferior parietal lobule | 40 | CON, FPN | -60 | -48 | 41 | 641 |
| b. Meta-analysis, Stop >Go & No-Think >Think |  |  |  |  |  |  |  |  |
| 1 | Right | Inferior frontal gyrus (VLPFC)<br>Insula | 44, 45 | CON, FPN | 36 | 26 | 0 | 4523 |
| 2 | Right/Left | Supplementary motor area | 6, 8 | CON, FPN, LAN | 14 | 14 | 60 | 3071 |
| 3 | Left | Inferior frontal gyrus<br>Insula | 44, 45 | CON, FPN | -44 | 18 | 0 | 2970 |
| 4 | Right | Inferior parietal lobule | 40 | CON, FPN, PMM | 58 | -46 | 34 | 2633 |
| 5 | Right | Anterior cingulate cortex | 24, 32 | CON, FPN | 6 | 22 | 38 | 1620 |
| 6 | Right | Middle frontal gyrus (DLPFC)<br>Superior frontal gyrus (DLPFC) | 9, 10, 46 | CON | 36 | 50 | 22 | 844 |
| 7 | Right | Basal ganglia |  |  | 16 | 8 | 8 | 776 |
| 8 | Left | Inferior parietal lobule | 40 | CON, FPN | -60 | -50 | 34 | 608 |
| 9 | Right | Precentral gyrus | 6 | CON, LAN | 44 | 2 | 46 | 270 |
| 10 | Right | Superior parietal lobule | 7 | FPN, DAN | 34 | -48 | 46 | 176 |
| c. Within-subjects & Meta-analysis, Stop >Go & No-Think >Think |  |  |  |  |  |  |  |  |
| 1 | Right | Inferior frontal gyrus (VLPFC)<br>Insula | 44, 45 | CON, FPN | 45 | 18 | 8 | 2666 |
| 2 | Right | Inferior parietal lobule | 40 | CON, FPN, PMM | 63 | -42 | 38 | 1620 |
| 3 | Right | Supplementary motor area | 6, 8 | CON, FPN, LAN | 15 | 18 | 64 | 1418 |
| 4 | Right | Middle frontal gyrus (DLPFC) | 9, 10, 46 | CON | 33 | 39 | 26 | 338 |
| 5 | Left | Inferior parietal lobule | 40 | CON, FPN | -60 | -48 | 41 | 270 |
| 6 | Right | Precentral gyrus | 6 | CON, LAN | 42 | 3 | 41 | 135 |

**Figure 3.**
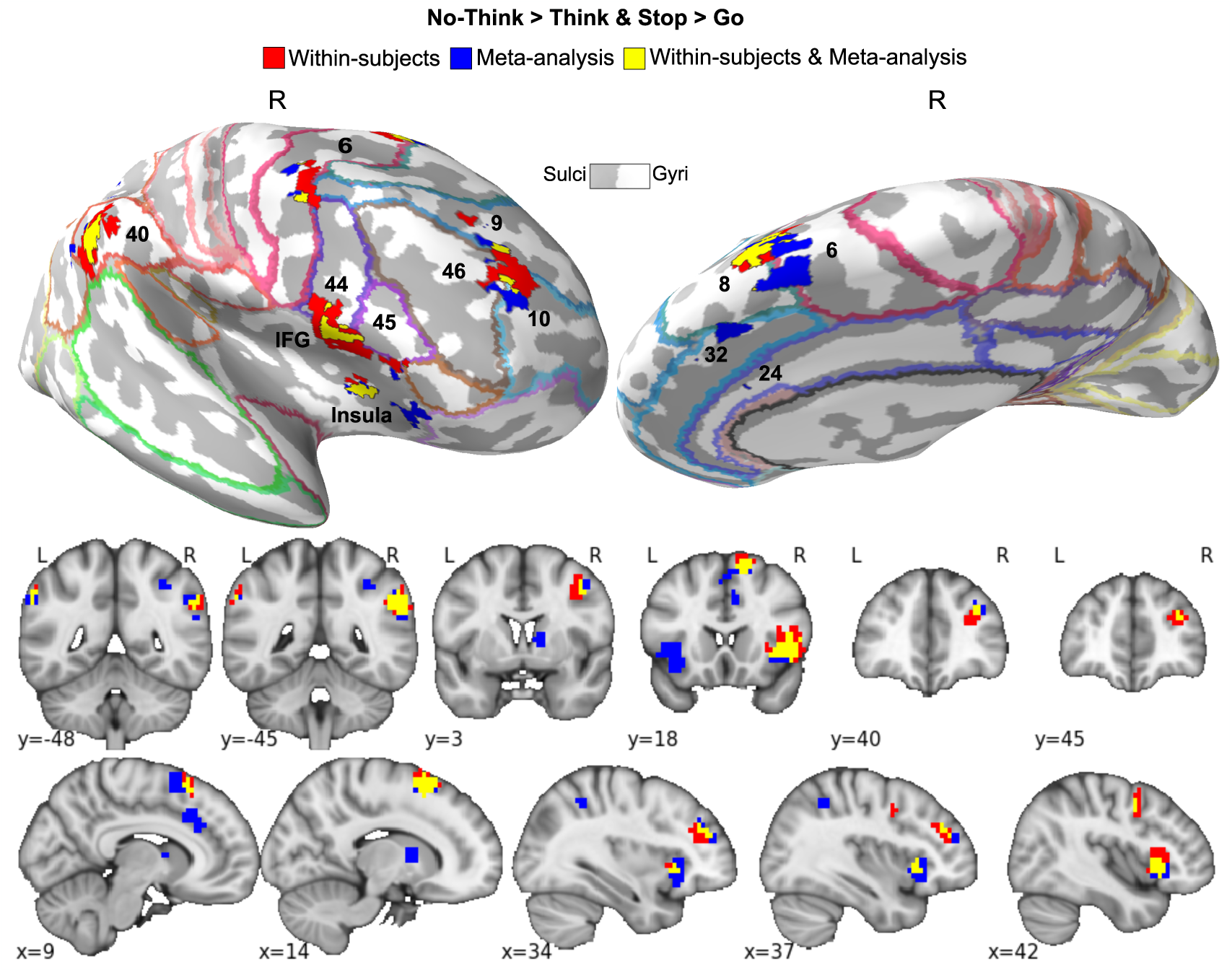
Domain-general inhibition-induced activations. Red: within-subjects (N = 24) conjunction of the Stop > Go and the No-Think > Think contrasts thresholded at p < 0.05 FDR corrected for whole-brain multiple comparisons. Blue: meta-analytic conjunction of Stop > Go and the No-Think > Think contrasts from independent 40 Stop-signal and 16 Think/No-Think studies. Yellow: overlap of the within-subjects and meta-analytic conjunctions. Results are displayed on an inflated MNI-152 surface with outlined and numbered Brodmann areas (top panel), as well as on MNI-152 volume slices (bottom panel). The brain images were generated using FreeSurfer software (http://surfer.nmr.mgh.harvard.edu), and PySurfer (https://pysurfer.github.io) and Nilearn (https://nilearn.github.io) Python (Python Software Foundation, DE, USA) packages.

**Figure 4.**
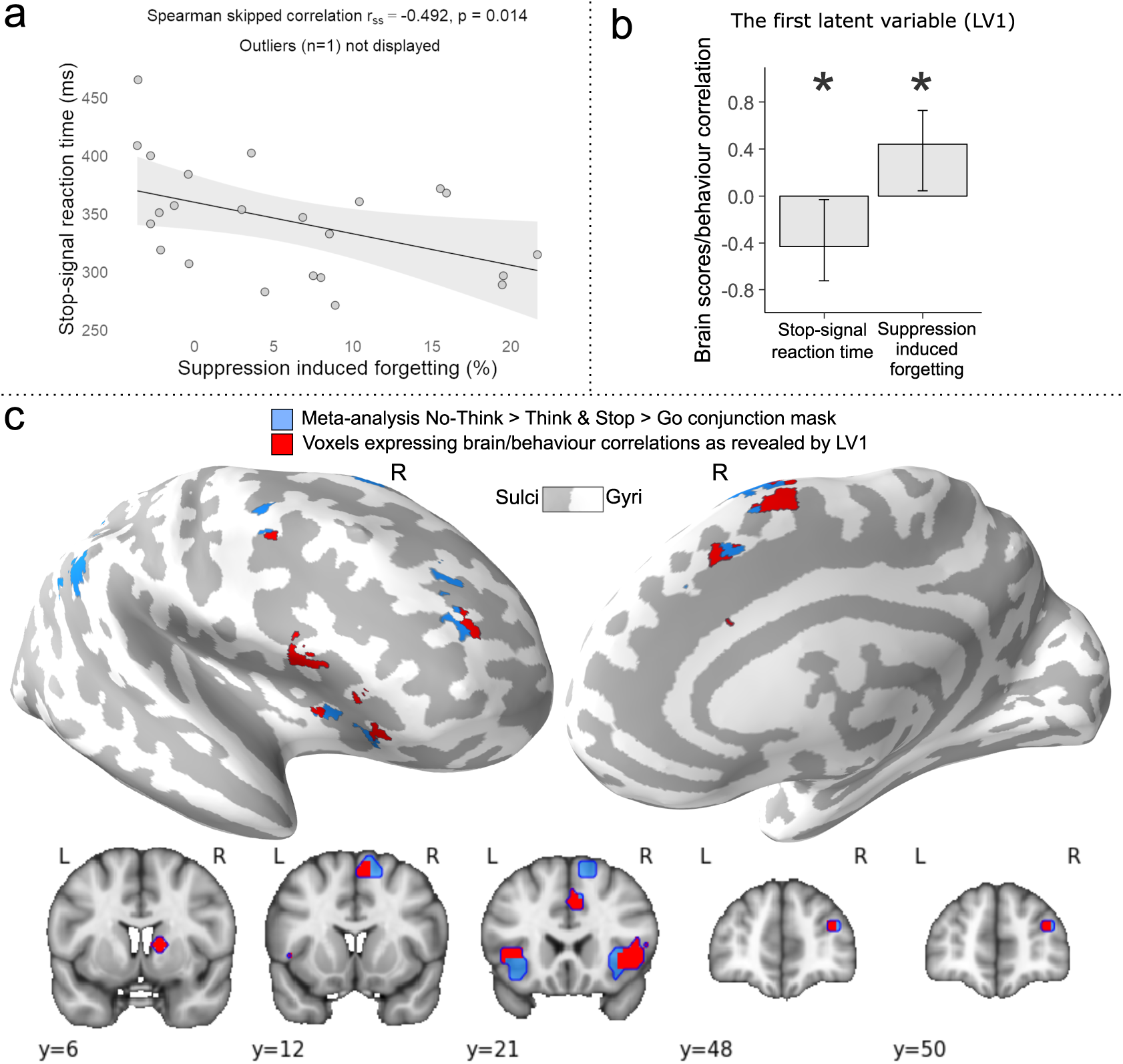
Domain-general behavioural and brain/behaviour relationships. **(a)** Better action stopping efficiency (shorter stop-signal reaction time) was associated with better inhibitory control over thoughts (percentage of items forgotten for No-Think relative to Baseline conditions at the final recall phase, i.e. suppression-induced forgetting; r_ss_ = −.492, p = .014). One bivariate outlier is not displayed on the scatterplot. Shading represents 95% CI. **(b and c)** A behavioural partial least squares (PLS) analysis was conducted to identify brain areas where individual variation in inhibition ability (SSRT and SIF) was related to increased inhibition-induced activity (main effect contrast of inhibition from the within-subject experiment, masked by the meta-analytic conjunction). **(b)** The first latent variable (LV1) identified voxels showing a significant pattern of brain/behaviour correlations to both SSRT and SIF (error bars indicate bootstrapped 95% CI), **(c)** The voxel salience map expressing LV1. Blue: meta-analytic conjunction mask. Red: voxels showing a significant pattern of brain/behaviour correlations as revealed by the LV1; thresholded at bootstrapped standard ratio 1.96, corresponding to p < 0.05, two-tailed. Results are displayed on an inflated MNI-152 surface (top panel), as well as on MNI-152 volume slices (bottom panel). The brain images were generated using FreeSurfer software (http://surfer.nmr.mgh.harvard.edu), and PySurfer (https://pysurfer.github.io) and Nilearn (https://nilearn.github.io) Python (Python Software Foundation, DE, USA) packages.

The observation that rDLPFC contributes to inhibitory control might seem surprising, given the published emphasis on the rVLPFC in motor inhibition studies (Aron et al., 2004, 2014). It could be that rDLPFC activation arises from the need to alternate between the Stop-signal and Think/No-Think tasks, or from carryover effects between tasks. We therefore compared the activations observed in our within-subjects conjunction analysis to a meta-analytic conjunction analysis of independent Stop-signal (N = 40) and Think/No-Think (N = 16) studies (see Methods) conducted in many different laboratories with different variations on the two procedures (see Guo et al., 2018) for an earlier version with fewer studies). The meta-analytic conjunction results were highly similar to our within-subjects results, with conjoint clusters in matched regions of DLPFC, VLPFC (BA44/45, extending into insula), right anterior cingulate cortex, and right basal ganglia (see Table 1b&c and Figure 3). Notably, in both the within-subjects and meta-analytic conjunctions, the domain-general activation in rDLPFC did not spread throughout the entire right middle frontal gyrus but was confined to the anterior portion of the rDLPFC, spanning BA9/46 and BA10. The convergence of these conjunction analyses suggests that the involvement of the rDLPFC, and our findings of conjoint activations across the two inhibitory domains more broadly, do not arise from the specific procedures of the inhibition tasks or to carryover effects arising from our within-subjects design; rather, they indicate a pattern that converges across laboratories and different experimental procedures.

The domain-general stopping activations included areas outside of the prefrontal cortex (see Table 1a and Figure 3). We characterised these activations in relation to large-scale brain networks, using a publicly available Cole-Anticevic brain-wide network partition (CAB-NP) (Ji et al., 2019). We used the Connectome Workbench software (Marcus et al., 2011) to overlay our activations over the CAB-NP to estimate the parcel and network locations of our clusters. Domain-general clusters primarily were located in the Cingulo-Opercular (CON) and Frontoparietal (FPN) networks (86% of parcels fell within these two networks in the within-subjects conjunction), but also included Posterior-Multimodal and Language networks parcels (see Table S1 and Figure S1). Of the 21 cortical parcels identified for the within-subjects conjunction (see Table S1), the majority (57%) participated in the CON, whereas 29% were involved in the FPN; the independent meta-analysis yielded similar findings (56% vs 30%; see Table S2 and Figure S2). Our main right prefrontal regions both featured parcels from the CON; however, whereas rDLPFC was located solely in the CON (in both the within-subjects and meta-analytic conjunctions), the rVLPFC region also included parcels from the FPN.

Together, these findings confirm the role of both the right anterior DLPFC and rVLPFC for both motor and memory inhibition. Moreover, they show that inhibitory control recruits a larger network of regions, dominated by the CON, and to a lesser degree, FPN. These findings suggest that domain-general inhibitory control may reflect a special configuration of the CON that includes elements of the FPN and other networks. Notably, key regions of the FPN were absent from all analyses, including the large middle frontal region often taken as a hallmark of domain-general cognitive control (Cole et al., 2013; Duncan, 2010).

### Right DLPFC and VLPFC support a common process underlying suppression-induced forgetting and action stopping efficiency

We next examined whether action inhibition and thought suppression depend on activity in the putative domain-general regions identified in our meta-analytic conjunction analysis. We tested whether activation in the very same voxels would predict SIF and SSRT. This test used behavioural PLS analysis (see Methods), excluding one behavioural bi-variate outlier from this analysis (see Methods), although the results with the outlier included did not qualitatively differ.

The first latent variable (LV) identified by PLS accounted for 78% of the covariance between inhibitory control activations and behavioural measures of SSRT and SIF. To specify how brain activation relates to those measures, we computed voxel saliences and a brain score for each participant (see Methods). A brain score indicates how much a participant expresses the multivariate spatial pattern of correlation between inhibitory control brain activations and behavioural measures of action and memory control captured by a LV. Thus, correlations between brain scores and behavioural measurements identify the direction and the strength of the relationship captured by a LV (i.e., the corresponding voxel salience over that LV). Within our meta-analytic conjunction regions (see Methods; Table 1b, Figure 3 and Figure 4c), participants’ brain scores for the first LV correlated negatively with SSRT scores (r = −0.432, [−0.724, −0.030] bootstrapped 95% CI) and positively with SIF scores (r = 0.441, [0.044, 0.729] bootstrapped 95% CI; Figure 4b). In other words, for voxels with high positive salience for this LV, a higher BOLD signal for the Inhibit > Respond contrast predicted faster SSRTs (i.e., better action stopping speed) and larger amounts of SIF (i.e., better memory inhibition). Voxels associated with significant positive salience arose across the entire set of domain-general conjunction regions except for the inferior parietal lobules (see Table 2 and Figure 4c). No voxels were associated with a significant negative salience (i.e., the opposite pattern).

**Table 2.** Control network regions showing a significant pattern of brain/behaviour correlations as revealed by the first latent variable of the PLS analysis.

| Brain region |  | ~BA | MNI of the peak |  |  | Volume (mm<sup>3</sup>) |
| --- | --- | --- | --- | --- | --- | --- |
|  |  |  | x | y | z |  |
| Right | Inferior frontal gyrus (VLPFC)<br>Insula | 44, 45 | 45 | 21 | 0 | 3375 |
| Right | Anterior cingulate cortex | 24, 32 | 6 | 30 | 34 | 1418 |
| Left | Inferior frontal gyrus<br>Insula | 44, 45 | -33 | 21 | 4 | 1046 |
| Right/Left | Supplementary motor area | 6, 8 | 6 | 9 | 64 | 1013 |
| Right | Basal ganglia |  | 15 | 3 | 8 | 709 |
| Right | Middle frontal gyrus (DLPFC) | 10, 46 | 33 | 48 | 19 | 304 |
| Right | Precentral gyrus | 6 | 42 | 3 | 41 | 68 |

These findings support the hypothesis that the stopping-evoked activity identified in our conjunction analyses plays behaviourally important roles both in stopping actions efficiently and in forgetting unwanted thoughts, a key attribute necessary to establish dynamic targeting.

### Stopping actions and stopping thoughts downregulates domain-specific target areas

A key attribute of dynamic targeting is that the domain-specific target areas are inhibited in response to activity of the domain-general source of inhibitory control, when the specific task goals require it. For example, inhibiting motor responses downregulates activity in M1 (Badry et al., 2009; Chowdhury et al., 2019; Mattia et al., 2012; Sumitash et al., 2019; Zandbelt & Vink, 2010), whereas inhibiting memory retrieval downregulates activity in the hippocampus (Anderson et al., 2016; Anderson & Hanslmayr, 2014; Anderson et al., 2004; Benoit & Anderson, 2012; Benoit et al., 2016; Benoit et al., 2015; Depue et al., 2007; Gagnepain et al., 2017; Hu et al., 2017; Levy & Anderson, 2012; Liu et al., 2016). Previously, we reported both of the foregoing patterns in a separate analysis of the current data (Schmitz et al., 2017). In the analyses below, we reconfirmed these findings using the left M1 and the right hippocampus ROIs which we defined specifically for the current DCM analyses (see Methods).

Dynamic targeting predicts a crossover interaction such that action stopping suppresses M1 more than it does the hippocampus, whereas thought stopping should do the reverse. A repeated-measures analysis of variance (ANOVA) confirmed a significant interaction between modulatory target regions (M1 vs. hippocampus) and stopping modality (stopping actions vs. stopping thoughts) on the BOLD signal difference between the respective inhibition and non-inhibition conditions in each modality (F_1,23_ = 42.71, p < 0.001; Figure 5a). Whereas stopping motor responses (Stop – Go) evoked greater downregulation of the M1 than the hippocampus ROI (t_23_ = 5.89, p < 0.001, d = 1.202), suppressing thoughts (No-Think – Think) evoked larger downregulation of the hippocampus than the M1 ROI (t_23_ = 3.22, p = 0.004, d = 0.658). Thus, action stopping and thought suppression preferentially modulated the left M1 and right hippocampus, respectively. Critically, these modulations were not solely produced by up-regulation in the Go or Think conditions, as illustrated by negative BOLD response during Stop (t_23_ = −3.88, p < 0.001, d = 0.791) and No-Think (t_23_ = −1.84, p = 0.04, d = 0.375) conditions (see Figure 5b). Thus, brain regions involved in representing the type of content requiring inhibition for each stopping task showed evidence of interrupted function during stopping, consistent with the requirements of dynamic targeting.

**Figure 5.**
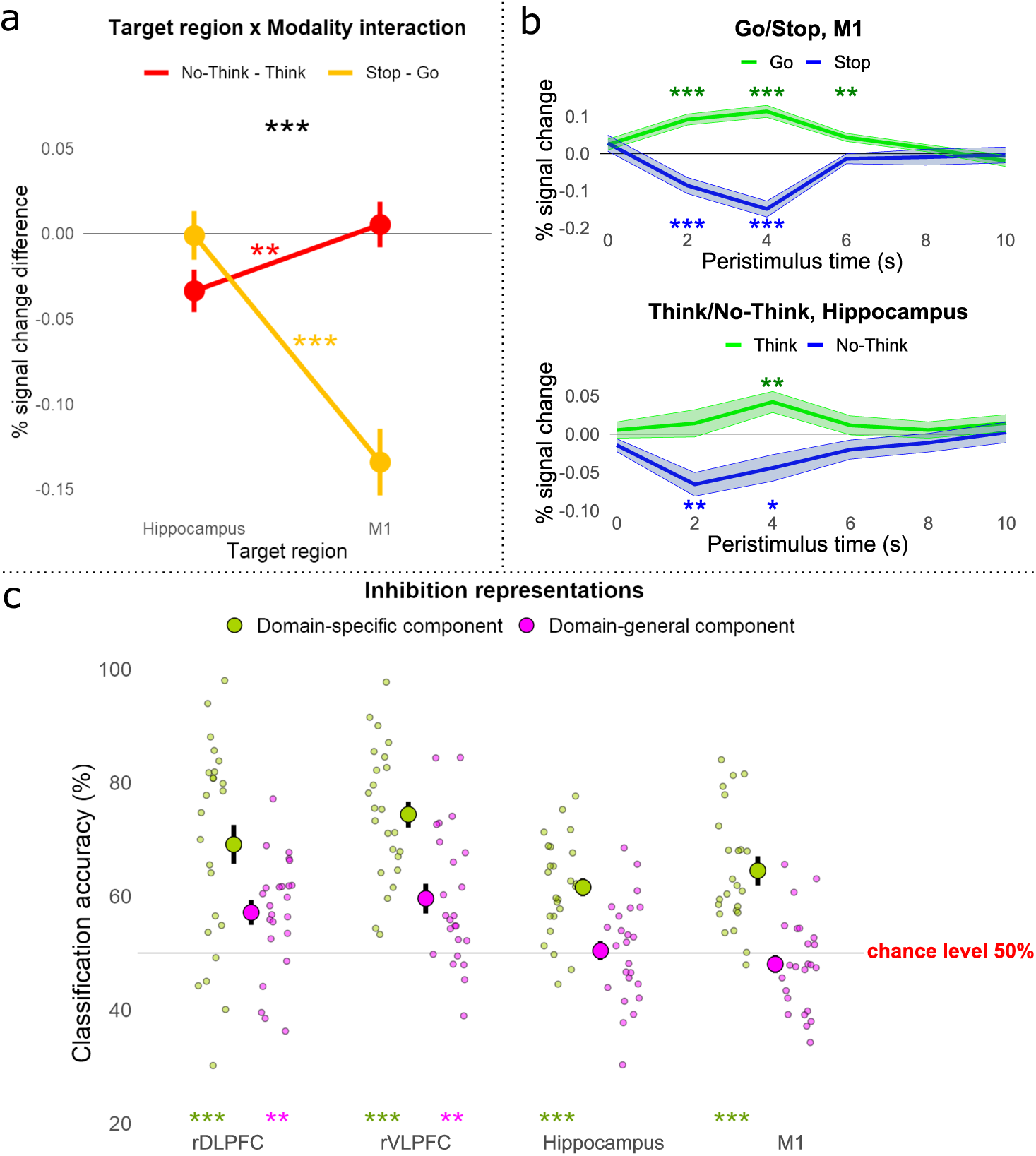
ROI analysis of domain-specific and domain-general modulation during thought and action suppression. ***p < 0.001; **p < 0.01; *p < 0.05. Error bars represent within-subject standard error. **(a)** Target areas M1 and hippocampus were modulated in a domain-specific manner. We calculated the BOLD signal in each target ROI for each condition by averaging across the time points from 2 to 8 s post-stimulus onset and subtracting out the onset value to account for pretrial variability. Then we subtracted the values of Go from Stop and Think from No-Think and entered them into a region by modality repeated-measures ANOVA. The ANOVA confirmed a significant interaction between modulatory target regions and stopping modality. Stopping actions (in yellow) evoked greater downregulation of M1 than of the hippocampus but suppressing thoughts (in red) evoked greater downregulation of the hippocampus than of M1. **(b)** The BOLD signal time-course in M1 (top panel) and hippocampus (bottom panel). During inhibition conditions (Stop and No-Think; in blue), the BOLD signal decreased below the baseline, whereas during respond conditions (Go and Think; in green) the BOLD signal increased above the baseline. **(c)** Using MVPA, we tested whether action and thought inhibition share a common voxel activation pattern within the four ROIs. We performed two types of pattern classification to identify domain-general (cross-task classification; in violet) and domain-specific (between-task classification; in green) components within each ROI. Large circles represent group average classification accuracies, and small circles represent individual participant accuracies.

### Action and thought stopping share common representations in the right DLPFC and VLPFC, but not in targeted regions

It is possible that despite the shared locus of activation in the rDLPFC and rVLPFC, the pattern of activation across voxels within these regions may fundamentally differ for action and thought stopping, a possibility that cannot be excluded with conventional univariate methods. However, dynamic targeting predicts similarities in the multivariate pattern of inhibitory control activity across voxels in the two tasks. Similarities should arise because of the shared engagement of a modality independent stopping process, even if some differences arise because of the stimulus processing and output pathways uniquely required to by each stopping process. To identify the similarities, we trained a classifier on the difference between Inhibit and Respond conditions in one modality and tested the ability to classify Inhibit and Respond conditions in the other domain. Such cross-modality decoding should not be possible in domain-specific target regions, reflecting their specialised involvement in action or memory stopping.

We performed the classification analysis on the rDLPFC, rVLPFC, right hippocampus, and left M1 ROIs which we defined for our DCM analyses (see Methods). The cross-modality classification revealed that a classifier trained on one modality could discriminate Inhibition from Respond conditions in the other modality significantly above chance (50%) for both rDLPFC (M = 57%, SD = 10%, one-tailed t_23_ = 3.48, p = 0.004, d = 0.711) and rVLPFC (M = 60%, SD = 12%, one-tailed t_23_ = 3.93, p = 0.001, d = 0.802). This cross-task decoding suggests a domain-general inhibitory control process in these regions (see Figure 5c). We also sought to identify differences in the patterns of activation across tasks by training a classifier to discriminate Stop from No-Think trials (see Methods). We found a significant domain-specific component in both rDLPFC (M = 69%, SD = 18%, one-tailed t_23_ = 5.09, p < 0.001, d = 1.039) and rVLPFC (M = 74%, SD = 12%, t_23_ = 10.10, p < 0.001, d = 2.06).

In contrast to the patterns observed in the prefrontal cortex, we observed no evidence of cross-task decoding in the modality-specific regions targeted by inhibitory control. This pattern arose for both right hippocampus (M = 50%, SD = 9%, one-tailed t_23_ = 0.23, p = 1, d = 0.046) and also left M1 (M = 48%, SD = 8, one-tailed t_23_ = −1.15, p = 1, d = −0.235), in which the cross-modality classifier accuracy did not significantly differ from chance performance (see Figure 5c). Nevertheless, these putative target regions responded very differently to the two modalities of inhibitory control, as evidenced by presence of significant domain-specific information in each region. A classifier could reliably distinguish No-Think trials from Stop trials within both the right hippocampus (M = 62%, SD = 9%, t_23_ = 6.59, p < 0.001, d = 1.346) and left M1 (M = 65%, SD = 10%, t_23_ = 6.85, p < 0.001, d = 1.399; see Figure 5c).

Because we z-normalised activation within each of these regions within each task, the ability to distinguish No-Think from Stop trials was not based on differences in overall univariate signal, but instead on information contained in distinct patterns of activity in each task. These findings reinforce the assumption that the hippocampus and M1 are uniquely affected by thought and action stopping respectively, as expected for domain-specific targets of inhibitory control. Taken together, these contrasting findings from the PFC and domain-specific regions are compatible with the view that rDLPFC and rVLPFC jointly contribute to a domain-general stopping process that dynamically targets different regions, depending on the nature of the content to be suppressed.

### Adaptive forgetting can be predicted using action stopping representations

Because dynamic targeting posits that LPFC contains domain-general stopping representations, training a classifier to distinguish stopping in one domain should predict stopping behaviour in other domains. For example, the ability of an action stopping classifier to distinguish when people are suppressing thoughts raises the intriguing possibility that it also may identify participants who successfully forget those thoughts. To test this possibility, we capitalised on an active forgetting phenomenon known as the conflict reduction benefit (for a review, see Anderson and Hulbert, 2021). The conflict-reduction benefit refers to the declining need to expend inhibitory control resources that arises when people repeatedly suppress the same intrusive thoughts. This benefit arises because inhibitory control induces forgetting of inhibited items, which thereafter cause fewer control problems. For example, over repeated inhibition trials, activation in rDPLFC, rVLPFC, and anterior cingulate cortex decline, with larger declines in participants who forget more of the memories they suppressed (Anderson & Hulbert, 2021; Kuhl et al., 2007; Wimber et al., 2015). If an action stopping classifier detects the inhibition process, two findings related to conflict-reduction benefits should emerge. First, over Think/No-Think task blocks, the action-stopping classifier should discriminate thought suppression less well, with high classification in early blocks that drops as memories are inhibited. Second, this decline should be larger for people showing greater SIF.

We examined how accurately an action stopping classifier distinguishes No-Think from Think conditions for the 8 fMRI runs. The rDLPFC showed a robust linear decline (F_7,157_ = 11.19, p = 0.001) in classification accuracy from the first (M = 77%) to the eighth (M = 40%) run, consistent with a conflict-reduction benefit (see Figure S4A). The rVLPFC exhibited a marginal linear decline (F_1,157_ = 3.04, p = 0.083) in classification accuracy from the first (M = 64%) to the eighth (M = 32%) run (see Figure S5A). Critically, for both rDLPFC (r_ss_ = −0.618, p = 0.001; Figure S4B) and rVLPFC (r_ss_ = −0.682, p < 0.001; Figure S5B), participants showing greater SIF exhibited a steeper classification accuracy decline. This suggests that adaptive forgetting had diminished demands on inhibitory control. Consistent with the involvement of inhibition, the decline in classifier performance also was associated to SSRT for both rDLPFC (r = 0.525, p = 0.008; Figure S4C) and rVLPFC (r_ss_ = 0.590, p = 0.002; Figure S5C). These findings support the view that suppressing unwanted thoughts engages a domain-general inhibition process indexed by action stopping and suggests that both rDLPFC and rVLPFC support this process.

### Right DLPFC and VLPFC dynamically couple with their domain-specific target areas to down-regulate their activity

Although rDLPFC and rVLPFC contribute to action and thought stopping, it remains to be shown whether either or both regions causally modulate target regions during each task, one of the five key attributes of dynamic targeting. On the one hand, rVLPFC alone might show dynamic targeting, exerting inhibitory modulation on the hippocampus or M1 in a task-dependent manner, as emphasized in research on motor response inhibition (Aron et al., 2004, 2014); rDLPFC may only be involved to maintain the inhibition task set in working memory, possibly exerting a modulatory influence on rVLPFC to achieve this (rVLPFC alone model). On the other hand, rDLPFC alone might show dynamic inhibitory targeting, consistent with the emphasis on the rDLPFC as the primary source of inhibitory control in research on thought suppression (Anderson & Hanslmayr, 2014; Anderson & Hulbert, 2021); rVLPFC may only be involved when attention is captured by salient stimuli, such as the stop signal or intrusions, possibly exerting a modulatory effect on rDLPFC to upregulate its activity (rDLPFC alone model). A third possibility is that rDLPFC and rVLPFC each contribute to top-down modulation in a content-specific manner, with only rDLPFC modulating the hippocampus during memory control, but only rVLPFC modulating M1 during action stopping. By this independent pathway hypothesis, both structures are pivotal to inhibitory control functions, but only with respect to their special domains, contrary to dynamic targeting. Finally, both rDLPFC and rVLPFC may be involved in dynamic targeting, modulating both hippocampus and M1 in a task-dependent manner; they may interact with one another to support stopping (Parallel modulation hypothesis).

To determine the way that rDLPFC and rVLPFC interact with each other and with the target regions of inhibitory control (M1 and hippocampus) we analysed effective connectivity between regions using dynamic causal modelling (DCM, see Methods). DCM accommodates the polysynaptic mediation of the causal influence that prefrontal regions could exert on activity in the hippocampus and in M1 (Anderson et al., 2016). DCM is ideally suited to test our hypotheses about which prefrontal regions drive inhibitory interactions, whether these vary by task context, and whether and how those prefrontal regions interact with one another to achieve inhibitory control.

Our model space included a null model with no modulatory connections and 72 distinct modulatory models (see Figure 6a) differing according to whether the source-target modulation was bidirectional, top-down, or bottom-up, whether rDLPFC, rVLPFC or both were sources of modulation, whether rDLPFC and rVLPFC interacted during inhibition tasks, and whether the site on which top-down modulation acted was appropriate to the inhibition task or not. We first compared the null model and models in which the direction of source-target modulation was either bidirectional, top-down, or bottom-up (24 models in each of the three families). The findings from these connectivity analyses were unambiguous. Bayesian Model Selection (BMS) overwhelmingly favoured models with bidirectional connections between the sources (rDLPFC and rVLPFC) and targets (M1 and hippocampus) with an exceedance probability (EP) of 0.9999. In contrast, the null modulation, top-down, and bottom-up models had EP of 0/0.0001/0, respectively (see Figure 6b). Exceedance probability refers to the extent to which a model is more likely in relation to other models considered. The bidirectional modulation confirms the existence of a top-down (our focus of interest) influence that prefrontal regions exert on activity in the hippocampus and in M1, alongside bottom-up modulation.

**Figure 6.**
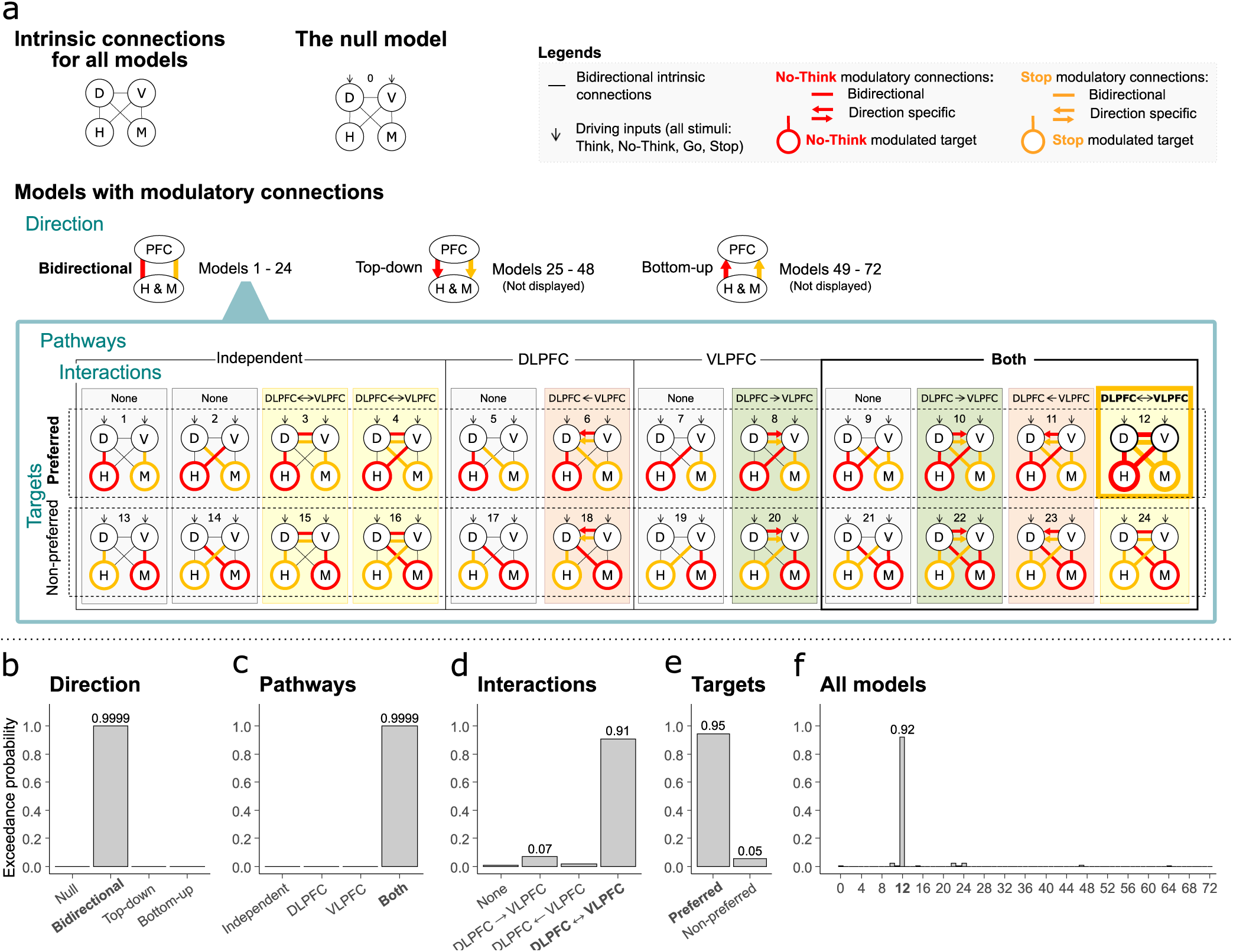
DCM model space and results. **(a)** DCM analysis determined the most likely inhibition-related interactions between domain-general inhibitory control source areas (D: rDLPFC, V: rVLPFC) and domain-specific target areas (H: right hippocampus, M: left M1). We compared 73 alternative models grouped into four family types. Direction: three families according to whether the source-target modulation is bidirectional, top-down, or bottom-up (we display only the 24 models within the bidirectional family as the further grouping was identical within each of the three families). Pathways: four families differing according to how Stop and No-Think modulate the pathways: independent modulation of target regions by rDLPFC and rVLPFC; rDLPFC only modulation; rVLPFC only modulation; or modulation by both rDLPFC and rVLPFC. Interactions: four families differing according to how Stop and No-Think modulate interactions between the rDLPFC and rVLPFC regions: no interactions; rVLPFC modulates rDLPFC; rDLPFC modulates rVLPFC; or bidirectional interaction between rDLPFC and rVLPFC. Targets: two families differing according to whether Stop and No-Think modulate the prefrontal connectivity with the preferred targets (M1 when stopping actions and hippocampus when stopping thoughts) or with the non-preferred targets (hippocampus when stopping actions and M1 when stopping thoughts). BMS (reporting exceedance probability to which a model is more likely to other models considered) overwhelmingly favoured models with **(b)** bidirectional source-target modulation; **(c)** both rDLPFC and rVLPFC modulating both the hippocampus and M1; **(d)** bidirectional interactions between the rDLPFC and rVLPFC; **(e)** the preferred target modulation. **(f)** The overall winning model also was strongly favoured by BMS even when directly assessing all 73 models, side by side, without grouping them into model families.

We next compared, within the 24 bidirectional models (models 1-24, see Figure 6a), whether either rDLPFC or rVLPFC was the sole dominant top-down source of inhibitory control (rDLPFC only vs rVLPFC only models) to models in which both regions comprised independent modulatory pathways (independent pathways model) or instead, contributed cooperatively to achieving top-down inhibitory control (parallel inhibition model). The BMS overwhelmingly favoured models in which both rDLPFC and rVLPFC contributed to modulating both the hippocampus and M1 with an exceedance probability (EP) of 0.9999; in contrast, Independent Pathways, rDLPFC alone, and rVLPFC alone models had an EP of 0.0001/0/0, respectively (see Figure 6c).

We next sought to distinguish subfamilies within this parallel model (models 9-12, and 21-24, see Figure 6a) that varied according to whether and how rDLPFC and rVLPFC interacted during inhibition: No-interaction at all between rDLPFC and rVLPFC (none); Unidirectional interaction from rVLPFC to rDLPFC (unidirectional rVLPFC); Unidirectional interaction from rDLPFC to rVLPFC (unidirectional rDLPFC) and bidirectional interaction (rDLPFC and rVLPFC interact with each other). If rDLPFC and rVLPFC work as a functional unit to achieve inhibitory control, one would expect clear evidence that some form of interaction occurs. Consistent with this view, BMS strongly favoured models with bidirectional interactions between the rDLPFC and rVLPFC (EP = 0.91; EP for the none, unidirectional rDLPFC, and unidirectional rVLPFC being 0.01/0.07/0.02; see Figure 6d).

Next, we tested whether inhibitory control is dynamically targeted to the appropriate target structure (e.g., hippocampus or M1), depending on which process needs to be stopped (memory retrieval or action production). According to our hypothesis, the rDLPFC and rVLPFC should down-regulate hippocampal activity during thought suppression, but should instead modulate M1, during action stopping. To test this dynamic targeting hypothesis, we compared the two remaining models (12 and 24, see Figure 6a) within our winning parallel/bidirectional subfamily. In the “preferred targets” model, rDLPFC and rVLPFC modulated the hippocampus during thought suppression, but M1 during action stopping; in the “non-preferred targets” model, these structures modulated content-inappropriate targets (e.g. M1 during thought suppression, but hippocampus during action stopping). BMS strongly favoured the model with preferred (EP = 0.95) over the non-preferred (EP = 0.05) target modulation (see Figure 6e). Indeed, the overall winning model also was strongly favoured by BMS even when directly assessing all 73 models, side by side, without grouping them into model families and subfamilies (BMS = 0.92; see Figure 6f).

The preferential modulations of hippocampus or M1, depending on whether thoughts or actions are to be suppressed, confirm our key hypothesis that top-down modulation by rDLPFC and rVLPFC is dynamically targeted depending on participants’ task goals. Together, the results of the DCM analysis suggest that, when inhibiting a prepotent response, the domain-general inhibitory control regions, rDLPFC and rVLPFC, interact with each other and are both selectively coupled with M1 when stopping actions and selectively coupled with the hippocampus when stopping thoughts.

## Discussion

The current findings identify two regions within the right LPFC that possess a dynamic targeting capability supporting the inhibition of both unwanted motor actions and thoughts: anterior rDLPFC and rVLPFC. These regions exhibited the five attributes needed to infer dynamic targeting. Both are engaged by diverse domains of inhibitory control, a finding supported not only by a within-subject conjunction analysis, but also via a meta-analytical conjunction; both show evidence of cross-task decoding, indicating that the representations formed in these regions are sufficiently general so that they recur in highly different stopping domains. Both regions are relevant to individual variation in inhibitory efficiency in both action stopping and thought suppression. Indeed, the multivariate activation pattern for action stopping resembled that for thought suppression enough so that it could be used as a proxy to predict how successfully people had suppressed their thoughts. Both regions are engaged alongside significant down-regulations in domain-specific target regions that we predicted *a priori* likely would require top-down inhibition; and both prefrontal regions show top-down effective connectivity with M1 and hippocampus during action stopping and thought suppression, supporting a causal role in their down-regulation. Critically, effective connectivity from both rDLPFC and rVLPFC to these two target regions dynamically shifted as participants moved between action to thought stopping, as would be required of a domain-general mechanism that can be flexibly targeted to suppress specialised content in multiple domains.

Based on these and related findings, we propose that anterior rDLPFC and rVLPFC constitute key hubs for a domain-general inhibitory control mechanism that can be dynamically targeted at diverse content represented throughout the brain. We focused here on the stopping of simple manual actions and verbal thoughts. Given this approach, this study does not address the breadth of thought content that can be targeted by this mechanism. However, when considered alongside the growing literature on retrieval suppression, the breadth of content is considerable. For example, the anterior rDLPFC and rVLPFC regions identified in the meta-analytic conjunction have been observed during the suppression of a range of stimuli, including words (Anderson et al., 2004; Benoit & Anderson, 2012; Levy & Anderson, 2012), visual objects (Gagnepain et al., 2014; Mary et al., 2020), neutral and aversive scenes (Benoit et al., 2015; Depue et al., 2007; Gagnepain et al., 2017; Liu et al., 2016) and person-specific fears about the future (Benoit et al., 2016). In addition, during retrieval suppression, these frontal regions exert top-down inhibitory modulation not only of the hippocampus (Anderson et al., 2016; Levy & Anderson, 2012), but also of other domain-specific content regions, including areas involved in representing visual objects (Gagnepain et al., 2014; Mary et al., 2020), places (Benoit et al., 2015; Gagnepain et al., 2017), and also emotional content in the amygdala (Depue et al., 2007; Gagnepain et al., 2017). Content-specific modulations are triggered especially when these types of content intrude into awareness in response to a cue and need to be purged (Gagnepain et al., 2017), indicating that inhibition can be dynamically targeted to diverse cortical sites to meet control demands. The current findings broaden the scope of this mechanism further by showing that it is not limited to stopping retrieval processes, but also extends to stopping the preparation and execution of motor responses, consistent with a broad mechanism involved in self-control over action and thought.

We considered the possibility that one of these two prefrontal regions is central to implementing top-down inhibitory control, with the other providing upstream inputs essential to initiate successful inhibitory control. Our effective connectivity analysis probed alternative hypotheses about the way rDLPFC and rVLPFC interact during inhibitory control. RDLPFC might implement the true inhibitory signal, receiving salience detection input from rVLPFC that up-regulates rDLPFC function. Alternatively, rVLPFC may implement inhibition, with rDLPFC preserving task set by sending driving inputs to the rVLPFC. Our findings indicate that both structures contributed in parallel to top-down inhibitory control and interacted bidirectionally during both action and thought stopping. Little evidence suggested a strong asymmetry in how rDLPFC and rVLPFC interacted, as should arise if one region simply served a role in salience detection or task-set maintenance. These findings suggest that rDLPFC and rVLPFC act together to implement top-down inhibitory control. Although it might seem surprising that two spatially segregated prefrontal regions would act in concert to achieve this function, it seems less unusual considering their potential role in the Cingulo-Opercular network (CON). The majority of the regions identified in our inhibition conjunction analysis participate in this network, suggesting that it may play an important role in achieving inhibitory control. Given the strong integrated activity of this network, elements of which are distributed throughout the brain (Cocuzza et al., 2020; Cole et al., 2013), this suggests future work should examine how rDLPFC and rVLPFC work together with other elements of this network to achieve successful inhibitory control.

The current proposal contrasts with models that emphasise the primacy of either rVLPFC or rDLPFC in inhibitory control, and which have not addressed dynamic targeting to diverse content. Research on motor inhibition has emphasised the rVLPFC as the source of top-down inhibitory control (Aron et al., 2004, 2014), although without evidence to exclude the role of rDLPFC. Indeed, studies cited as favouring the selective role of rVLPFC often support contributions of the anterior rDLPFC structure identified here. For example, whereas intracranial stimulation in primates establishes the causal necessity of the rVLPFC in motor stopping, so too does stimulation of the dorsal bank of the principal sulcus, the putative monkey homologue of the rDLPFC in humans (Sasaki et al., 1989); and whereas intracranial recordings in humans show stopping-related activity in rVLPFC, they also reveal it in anterior rDLPFC and often prior to rVLPFC (Swann et al., 2013). Research on thought suppression has emphasised the rDLPFC as the source of top-down inhibitory control (Anderson et al., 2016; Anderson & Hanslmayr, 2014; Anderson et al., 2004); but most studies supporting the role of rDLPFC in thought suppression also reveal activations in the rVLPFC (Guo et al., 2018). Indeed, as our within-subjects and meta-analytic conjunctions unambiguously confirm, both regions are recruited during both inhibitory control tasks. The current study goes further than establishing conjoint activation: Pattern classification and connectivity analyses show the involvement of both regions in the dynamics of control, without selectivity. These findings validate the importance of both regions, establish the domain-generality of their influence, and demonstrate the dynamic inhibitory targeting capacity necessary to infer a flexible control mechanism.

The present findings highlight a potentially important difference between the brain networks involved in inhibitory control and other forms of cognitive control that do not require the inhibition of a motor or cognitive process. Maintaining rules in working memory, implementing task sets, performing multi-tasking, and manipulating information actively are all clear cases of cognitive control that can require interference resolution, but do not necessarily entail active stopping. The above tasks engage the widely discussed fronto-parietal network (FPN), often assigned a central role in implementing cognitive control more broadly (Cole et al., 2013; Cole & Schneider, 2007; Duncan, 2010; Fox et al., 2005). One might assume that because inhibitory control is a form of cognitive control that the FPN would be central to it as well. Nevertheless, the FPN, though involved in our tasks, appeared less prominent than the CON, which accounted for the majority of distinct cortical parcels participating in our domain-general inhibition regions. We found little evidence for involvement of major areas of the FPN, including much of the middle frontal gyrus bilaterally in our multimodal inhibition regions. As our meta-analysis and within-subjects comparisons confirm, inhibitory control is strongly right lateralised, which also is not a feature emphasised in research on the FPN. Our findings raise the possibility that stopping actions and thoughts may rely on a distinct network, with different functional characteristics to the FPN.

Dynamic inhibitory targeting provides a neurocognitive framework that can account for both associations and dissociations in the abilities to suppress unwanted thoughts and actions. On the one hand, deficits in both action and thought stopping should arise with dysfunction in the rDLPFC or rVLPFC, given the common reliance of these abilities on those regions. Such associations occur frequently. In the general population, people scoring highly on self-report measures of impulsivity or compulsivity also report greater difficulty with intrusive thoughts (Gay et al., 2011; Gillan et al., 2016). Clinically, persistent intrusive thoughts and action stopping deficits co-occur in numerous disorders: Obsessive thoughts and compulsive actions in obsessive-compulsive disorder (Fineberg et al., 2018; Gillan et al., 2017); intrusive memories and impaired response inhibition in PTSD (Falconer et al., 2008; Sadeh et al., 2018; Sadeh et al., 2015; van Rooij & Jovanovic, 2019; Wu et al., 2015); persistent worry and impulsivity in anxiety disorders (Berg et al., 2015) and intrusive thoughts and compulsivity in addiction (Everitt & Robbins, 2016; Kavanagh et al., 2005; May et al., 2015). These co-morbid deficits may reflect dysfunction in the rDLPFC, the rVLPFC or in other shared components of their control pathways. On the other hand, dissociations should arise when dysfunction selectively disrupts a domain-specific pathway linking rLPFC to target sites involved in generating actions and thoughts, including dysfunction to local inhibition at the target site itself. For example, individual variation in local GABAergic inhibition within the hippocampus or M1 predict inhibitory control over memories and actions, respectively, independently of prefrontal function (He et al., 2019; Schmitz et al., 2017). Thus, selective difficulties in action stopping or thought inhibition may arise, given focal deficits in either motor cortical or hippocampal GABA (Schmitz et al., 2017). The separate contributions of domain-general and domain-specific factors to inhibitory control implied by dynamic targeting constrains the utility of motor inhibition as a metric of inhibitory control over thought and may explain the surprisingly small SSRT deficits in major depression and anxiety, relative to attention deficit hyperactivity disorder or obsessive-compulsive disorder (Lipszyc & Schachar, 2010).

The current study did not seek to characterise the polysynaptic pathways through which the rDLPFC and rVLPFC suppress activity in either M1 or the hippocampus (Anderson et al., 2016; Depue et al., 2016). Rather, we focused on the existence of a central, domain-general inhibitory control function capable of flexibly shifting its top-down influence across actions and thoughts. By juxtaposing two well characterised model systems for stopping actions and thoughts, each with distinct neural targets of inhibition, we were able to show that the same set of prefrontal regions is involved in stopping processing in different cortical target areas, in a rapid, flexible manner. In doing so, we established evidence for dynamic inhibitory targeting as a key mechanism of domain-general inhibitory control in the human brain. More broadly, this work suggests that the human capacity for self-control in the face of life’s challenges may emerge from a common wellspring of control over our actions and thoughts.

## Methods

We used a dataset from a published study (Schmitz et al., 2017). However, here all data were independently re-analysed with a different focus.

### Participants

Thirty right-handed native English speakers participated. Participants gave written informed consent and received money for participating. Five participants did not reach the 40% learning criterion on the Think/No-Think task, and one fell asleep during fMRI acquisition. The final sample comprised 24 participants (7 males, 17 females), 19-36 years old (M = 24.67 years, SD = 4.31). Participants had normal or corrected-to-normal vision and no reported history of neurological, medical, or memory disorders, and they were asked not to consume psychostimulants, drugs, or alcohol before the experiment. The Cambridge Psychology Research Ethics Committee approved the project.

### Experimental paradigm

Participants performed adapted versions of the Stop-signal (Logan & Cowan, 1984) and Think/No-Think (Anderson & Green, 2001) tasks. Both tasks require participants to stop unwanted processes, but in the motor and memory domains, respectively.

The Stop-signal task assesses the ability to stop unwanted actions. Participants first learn stimulus-response associations and then perform speeded motor responses to the presented (Go) stimuli. Occasionally, shortly after the Go stimulus, a stop signal occurs, and participants must withhold their response. We measured the stop-signal reaction time (SSRT), an estimate of how long it takes the participant to stop.

The Think/No-Think task assesses the ability to stop unwanted memory retrievals. Participants first form associations between unrelated cue-target word pairs. Then participants receive two-thirds of the cues as reminders (one at a time) and are asked to either think (Think items) or to not-think (No-Think items) of the associated target memory, with each Think and No-Think reminder repeated numerous times throughout the task. Finally, participants attempt to recall all initially learned associations. Typically, recall performance suffers for No-Think items compared to Baseline items that were neither retrieved nor suppressed during the think/no-think phase. This phenomenon, known as suppression-induced forgetting (SIF), indirectly measures the ability to stop unwanted memory retrievals by quantifying inhibitory aftereffects of this process (Anderson & Hanslmayr, 2014; Anderson & Weaver, 2009).

### Stimuli and apparatus

We presented stimuli and recorded responses with Presentation software (Neurobehavioral Systems, Albany, CA, USA). For the Stop-signal task, four visually discriminable red, green, blue, and yellow coloured circles of 2.5 cm in diameter, presented on a grey background, constituted the Go stimuli (Figure 2a). Participants responded by pressing one of the two buttons (left or right) with a dominant (right) hand on a button box. An auditory 1000 Hz “beep” tone presented at a comfortable volume for 100 ms signalled participants to stop their responses. A fixation cross appeared in 50-point black Arial Rounded font on a grey background prior to the onset of the Go stimulus.

For the Think/No-Think task, we constructed 78 weakly relatable English word pairs (cue-target words, e.g., Part-Bowl) as stimuli and an additional 68 semantically related cue words for 68 of the target words (e.g., a cue word ‘Cornflake’ for the target word ‘Bowl’). We used 60 of the target words and their related and weak cues in the critical task, with the other items used as fillers. We divided the critical items into three lists composed of 20 targets and their corresponding weak cue words (the related word cues were set aside to be used as independent test cues on the final test; see procedure). We counterbalanced these lists across the within-subjects experimental conditions (Think, No-Think, and Baseline) so that across all participants, every pair participated equally often in each condition. We used the filler words both as practice items and also to minimise primacy and recency effects in the study list (Murdock, 1962). Words appeared in a 32-point Arial font in capital letters on a grey background (Figure 2b). During the initial encoding and final recall phases, we presented all cues and targets in black. For the Think/No-Think phase, we presented the Think cues in green and the No-Think cues in red, each preceded by a fixation cross in 50-point black Arial Rounded font on a grey background.

### Procedure

The procedure consisted of seven steps: 1) stimulus-response learning for the Stop-signal task: 2) Stop-signal task practice; 3) encoding phase of the Think/No-Think task; 4) Think/No-Think practice; 5) practice of interleaved Stop-signal and Think/No-Think tasks; 6) experimental phase during fMRI acquisition; 7) recall phase of the Think/No-Think task. We elaborate these steps below (see also Figure 2c).

#### Step 1 – Stop-signal task stimulus-response learning

Participants first formed stimulus-response associations for the Stop-signal task. As Go stimuli, we presented circles in four different colours (red, green, blue, and yellow) and participants had to respond by pressing one of the two buttons depending on the circle’s colour. Thus, each response button had two colours randomly assigned to it and participants associated each colour to its particular response.

Participants learned the colour-button mappings in two sets of two colours, with the first colour in a set associated with one button, and the second with the other button. After practising the responses to these colours in random order 10 times each, the same training was done on the second set. Subsequently, participants practised the colour-button mappings of all four colours in random order until they responded correctly to each colour on 10 consecutive trials. During the practice, we instructed participants to respond as quickly and accurately as possible and provided feedback for incorrect or slow (> 1000 ms) responses.

#### Step 2 – Stop-signal task practice

Once participants learned the stimulus-response associations, we introduced the Stop-signal task. We instructed participants to keep responding to each coloured circle as quickly and accurately as possible but indicated that on some trials, after the circle appeared, a beep would sound, and that they should not press any button on these trials. We also told participants to avoid slowing down and waiting for the beep, requesting instead that they treat failures to stop as normal and always keep responding quickly and accurately. Thus, on Go trials, participants responded as quickly as possible, whereas, on Stop trials, a tone succeeded the cue onset, signalling participants to suppress their response. To facilitate performance, participants received on-screen feedback for incorrect and too slow (> 700 ms) responses to Go trials, and for pressing a button on Stop trials.

Figure 2a presents the trial timings. Go trials started with a fixation cross, presented for 250 ms, followed by a coloured circle until response or for up to 2500 ms. After the response and a jittered inter-trial interval (M = 750 ms, SD = 158.7 ms), a new trial commenced. Stop trials proceeded identically except that a tone sounded shortly after the circle appeared. This stop signal delay varied dynamically in 50 ms steps (starting with 250 ms or 300 ms) according to a staircase tracking algorithm to achieve approximately a 50% success-to-stop rate for each participant. Note that the longer the stop signal delay is, the harder it is to not press the button. The dynamic tracking algorithm reduces participants’ ability to anticipate stop signal delay timing and provides a method for calculating the SSRT. In this practice step, participants performed 96 trials, of which 68 (71%) were Go trials and 28 (29%) were Stop trials.

#### Step 3 – Think/No-Think task encoding phase

Once participants had learned the Stop-signal task, we introduced the Think/No-Think task. In the encoding phase, participants formed associations between 60 critical weakly-related word pairs (e.g., Part-Bowl) and between 18 filler pairs. First, participants studied each cue-target word pair for 3.4 s with an inter-stimulus interval of 600 ms. Next, from each studied pair, participants saw the cue word only and recalled aloud the corresponding target. We presented each cue for up to 6 s or until a response was given. Six hundred ms after cue offset, regardless of whether the participant recalled the item, the correct target appeared for 1 s. We repeated this procedure until participants recalled at least 40% of the critical pairs (all but 5 participants succeeded within the maximum of three repetitions). Finally, to assess which word-pairs participants learned, each cue word appeared again for 3.3 s with an inter-stimulus interval of 1.1 s and participants recalled aloud the corresponding target. We provided no feedback on this test.

#### Step 4 – Think/No-Think practice

After participants encoded the word pairs, the Think/No-Think practice phase commenced. On each trial, a cue word appeared on the screen in either green or red. We instructed participants to recall and think of the target words for cues presented in green (Think condition) but to suppress the recall and avoid thinking of the target words for those cues presented in red (No-Think condition). Participants performed the direct suppression variant of the Think/No-Think task (Benoit & Anderson, 2012; Bergström et al., 2009) in which, after reading and comprehending the cue, they suppressed all thoughts of the associated memory without engaging in any distracting activity or thoughts. We asked participants to “push the memory out of mind” whenever it intruded.

Trial timings appear in Figure 2b. A trial consisted of presenting a cue in the centre of the screen for 3 s, followed by an inter-stimulus interval (0.5 s, M = 2.3 s, SD = 1.7 s) during which we displayed a fixation cross. We jittered the inter-stimulus interval (0.5 s, M = 2.3 s, SD = 1.7 s) to optimize the event-related design (as determined by optseq2: http://surfer.nmr.mgh.harvard.edu/optseq). In this practice phase, we used 12 filler items, six of which were allocated to the Think condition and six to the No-Think condition. We presented each item three times in random order (36 trials in total). In the middle of the practice, we administered a diagnostic questionnaire to ensure participants had understood and followed the instructions.

#### Step 5 – Interleaved Stop-signal and Think/No-Think practice

Before moving into the MRI scanner, participants performed an extended practice phase interleaving the Stop-signal and Think/No-Think tasks. For the Think/No-Think task, we again used 12 filler items. Other than that, and the fact that the practice took place outside the MRI scanner, this phase was identical to a single fMRI acquisition session described into more detail next.

#### Step 6 – Experimental phase and fMRI acquisition

In the main experimental phase, participants underwent 8 fMRI scanning runs in a single session. Before the scanning began, participants saw the correct button-colour mappings and all 78 word pairs briefly presented on the screen to remind them of the task and items. After the brief refresher, the fMRI acquisition started. During each fMRI run, participants performed 8 blocks of the Think/No-Think task interleaved with 8 blocks of the Stop-signal task. All blocks lasted 30 s. To minimize carry-over effects, we interspersed 4 s rest periods (blank screen with a grey background) between blocks. Each block began with items that we did not score (the filler items) to reduce task-set switching effects between blocks. Within each block, we pseudo-randomly ordered all trials, and the trial timings for both tasks were identical to those used in their respective practice phases (step 2 and step 4; Figure 2a Figure 2b).

Four of the Stop-signal task blocks contained Go trials only. We did not use these blocks in this report. Each of the other four Stop-signal blocks contained 12 trials, yielding 384 trials in total (8 runs * 4 blocks per run * 12 trials per block). On average, across participants, Stop trials constituted 32% (SD = 2%) of the trials. As in the practice phase, a staircase tracking algorithm varied the delay between cue onset and stop-signal tone according to each participant’s performance, keeping the stopping success at approximately 50%.

Each of the Think/No-Think blocks contained 6 trials, starting with a filler item as a Think trial followed by 5 Think or No-Think items in a pseudo-random order. Within each fMRI run, participants saw all 20 critical Think and 20 critical No-Think items once. Thus, across the 8 runs, participants recalled or suppressed each memory item 8 times. The proportion of the Think trials (58%) exceeded the proportion of the No-Think trials (42%) to better resemble the higher frequency of Go trials than Stop trials during the Stop-signal task. We accomplished this by assigning Think trials to the filler items, without changing the frequency of Think trials on critical experimental items. After the fourth (middle) run, to allow participants to rest, we acquired their anatomical scan and administered the diagnostic questionnaire to ensure that participants closely followed the instructions of the Think/No-Think task.

#### Step 7 – Think/No-Think recall phase

In the final step (inside the scanner but without any scan acquisition), we measured the aftereffects of memory retrieval and suppression via a cued-recall task on all word pairs (encoded in step 3). This included 20 Baseline items that were neither retrieved nor suppressed during the Think/No-Think phase and that thus provided a baseline estimate of the memorability of the pairs.

To reinstate the context of the initial encoding phase, we first tested participants on 10 filler cue words, 6 of which they had not seen since the encoding phase (step 3) and 4 of which they saw during the interleaved Stop-signal and Think/No-Think practice phase (step 5). We warned participants that the cues in this phase could be ones they had not seen for a long time and encouraged them to think back to the encoding phases to retrieve targets.

Following context reinstatement, participants performed the same-probe and independent-probe memory tests. In the same-probe test, we probed memory with the original cues (e.g. the weakly related cue word ‘Part’ for the target word ‘Bowl’). We included the independent-probe test to test whether forgetting generalized to novel cues (Anderson and Green, 2001), using the related cues we had designed for each target. For example, we cued with the semantic associate of the memory and its first letter (e.g., ‘Cornflake – B’ for the target ‘Bowl’). Across participants, we counterbalanced the order in which the tests appeared. In both tests, cues appeared for a maximum of 3.3 s or until participants gave a response, with an inter-stimulus interval of 1.1 s. We coded a response as correct if participants correctly recalled the target while the cue was onscreen.

Finally, we debriefed participants, and administered a post-experimental questionnaire to capture participants’ experiences and the strategies they used in the Think/No-Think and Stop-signal tasks.

### Brain image acquisition

We collected MRI data using a 3-Tesla Siemens Tim Trio MRI scanner (Siemens, Erlangen, Germany) fitted with a 32-channel head coil. Participants underwent eight functional runs of the blood-oxygenation-level-dependent (BOLD) signal acquisitions. We acquired functional brain volumes using a gradient-echo, T2*-weighted echoplanar pulse sequence (TR = 2000 ms, TE = 30 ms, flip angle = 90°, 32 axial slices, descending slice acquisition, voxel resolution = 3 mm^3^, 0.75 mm interslice gap). We discarded the first four volumes of each session to allow for magnetic field stabilisation. Due to technical problems encountered during task performance, we discarded from the analysis one functional run from two participants each, and two functional runs from another participant. After the fourth functional run, we acquired an anatomical reference for each participant, a high-resolution whole-brain 3D T1-weighted magnetization-prepared rapid gradient echo (MP-RAGE) image (TR = 2250 ms, TE = 2.99 ms, flip angle = 9°, field of view = 256 x 240 x 192 mm, voxel resolution = 1 mm^3^). Following the acquisition of the anatomical scan, participants underwent the remaining four functional runs.

### Data analysis

#### Behavioural performance

For statistical analyses of the behavioural data, we used R (v4, 2020-04-24) in Jupyter Notebook (Anaconda, Inc., Austin, Texas). The data and detailed analysis notebook are freely available at http://bit.do/analysis-domain-general. For all statistical comparisons, we adopted p < 0.05 as the significance threshold.

For correlation analyses, we followed recommendations by Pernet et al. (2013) and used one of three correlation methods depending on whether the data were normally distributed or contained outliers. If there were no outliers and data were normally distributed, we performed Pearson correlation and reported it as ‘r’. If there were univariate outliers (but no bivariate) or data were not normally distributed, we performed robust 20% Bend correlation and reported it as ‘r_pp_’. If there were bivariate outliers, we performed robust Spearman skipped correlation using the minimum covariance determinant (MCD) estimator and reported it as ‘r_ss_’. For univariate and bivariate outlier detection, we used boxplot and bagplot methods, respectively.

For the analysis of Stop-signal task data, we followed the guidelines by Verbruggen et al. (2019) and calculated SSRT using the integration method with the replacement of Go omissions. Specifically, we included all Stop trials and all Go trials (correct and incorrect), replacing missed Go responses with the maximum Go RT. To identify the nth fastest Go RT, we multiplied the number of total Go trials by the probability of responding to stop signal (unsuccessful stopping). The difference between the nth fastest Go RT and the mean SSD provided our estimate of SSRT.

In addition to SSRT, we calculated the probability of Go omissions, probability of choice errors on Go trials, probability of responding to Stop trials, mean SSD of all Stop trials, mean correct Go RT, and mean failed Stop RT. We also compared RTs of all Go trials against RTs of failed Stop trials to test the assumption of an independent race between a go and a stop runner. Besides, we assessed the change of Go RTs across the eight experimental blocks. Prior work suggests that the experiment-wide integration method can result in underestimation bias of SSRT if participants slow their RT gradually across experimental runs. In that case, a blocked integration method would provide a better measure of SSRT (Verbruggen et al., 2013). In our data, however, on average within the group, we observed a negligible decrease in RT across runs (B = −2.555, p = .250), suggesting that the experiment-wide integration method was more appropriate.

For the Think/No-Think task data, we focused on the critical measure: SIF. We used the final recall scores (from step 7) of No-Think and Baseline items conditionalized on correct initial training performance (at step 3), as in prior work (Anderson et al., 2004). Thus, in the final recall scores, we did not include items that were not correctly recalled (M = 29%, SD = 17) during the criterion test of the encoding phase, as the unlearned items can be neither suppressed nor retrieved during the Think/No-Think phase (step 6). As in our previous work (Schmitz et al., 2017), we averaged the scores across the same-probe and independent-probe tests and the difference between the Baseline and No-think item recall scores constituted our measure of SIF. To assess the group effect of SIF, we tested the data for normality (W = 0.95, p = 0.264) and performed a one-sample, one-sided t-test to determine if SIF is greater than zero. Finally, to assess whether inhibition ability generalises across motor and memory domains, we performed a correlation between the SSRT and SIF scores.

To identify univariate and bi-variate outliers in the SSRT and SIF scores, we used box plot method, which relies on the interquartile range. Univariate outliers were not present for any of the two measures. One bi-variate outlier was removed from the correlation analysis and the behavioural partial least squares analysis (described below). Nevertheless, outlier removal did not qualitatively alter the results.

#### Brain imaging data

##### Pre-processing

We pre-processed and analysed the brain imaging data using Statistical Parametric Mapping v12 release 7487 (SPM12; Wellcome Trust Centre for Neuroimaging, London) in MATLAB vR2012a (The MathWorks, MA, USA). To approximate the orientation of the standard Montreal Neurological Institute (MNI) coordinate space, we reoriented all acquired MRI images to the anterior-posterior commissure line and set the origins to the anterior commissure. Next, we applied our pre-processing procedure to correct for head movement between the scans (images realigned to the mean functional image) and to adjust for temporal differences between slice acquisitions (slice-time correction relative to the middle axial slice). The procedure then co-registered each participant’s anatomical image to the mean functional image and segmented it into grey matter, white matter, and cerebrospinal fluid. We then submitted the segmented images for each participant to the DARTEL procedure (Ashburner, 2007) to create a group-specific anatomical template which optimises inter-participant alignment. The DARTEL procedure alternates between computing a group template and warping an individual’s tissue probability maps into alignment with this template and ultimately creates an individual flow field of each participant. Subsequently, the procedure transformed the group template into MNI-152 space. Finally, we applied the MNI transformation and smoothing with an 8 mm full-width-at-half-maximum (FWHM) Gaussian kernel to the functional images for the whole-brain voxel-wise analysis.

##### Univariate whole-brain analysis

To identify brain areas engaged in both inhibiting actions and inhibiting memories, we performed a whole-brain voxel-wise univariate analysis. We high-pass filtered the time series of each voxel in the normalised and smoothed images with a cut-off frequency of 1/128 Hz, to remove low-frequency trends, and modelled for temporal autocorrelation across scans with the first-order autoregressive (AR(1)) process. We then submitted the pre-processed data of each participant to the first-level, subject-specific, General Linear Model (GLM) modelling a single design matrix for all functional runs.

We modelled the Stop-signal task and Think/No-Think task conditions as boxcar functions, convolved with a haemodynamic response function (HRF). In the model, we used group average response latencies for each trial type as the trial durations for the Stop-signal task condition, but we used 3 s epochs for the Think/No-Think task condition. As in the behavioural analysis, we conditionalized the Think and No-Think conditions on initial encoding performance. The main conditions of interest for our analysis included: correct Stop, correct Go (from the mixed Stop-signal and Go trial blocks only), conditionalized No-Think and conditionalized Think. Unlearned No-Think and Think items, filler items, incorrect Stop, incorrect Go and Go trials from the Go-only blocks we modelled as separate regressors of no interest. We also included the six realignment parameters for each run as additional regressors of no interest, to account for head motion artefacts, and a constant regressor for each run. We obtained the first-level contrast estimates for Stop, Go, No-Think, and Think conditions, and the main effect of Inhibit [Stop, No-Think] > Respond [Go, Think].

At the second-level random-effect group analysis we entered the first-level contrast estimates of Stop, Go, No-Think, and Think conditions into a repeated-measures analysis of variance (ANOVA), which used pooled error and correction for non-sphericity, with participants as between-subject factor. We then performed a conjunction analysis of Stop > Go No-Think > Think contrasts, using the minimum statistics analysis method implemented in SPM12, and testing the conjunction null hypothesis (Friston et al., 2005; Nichols et al., 2005). The results of the conjunction analysis represent voxels that were significant for each individual contrast thresholded at p < 0.05 false discovery rate (FDR) corrected for whole-brain multiple comparisons.

##### Behavioural partial least squares (PLS) analysis

We hypothesised that domain-general inhibitory control brain activity would be related to domain-general inhibitory behaviour. To test our hypothesis, we performed behavioural PLS analysis (Krishnan et al., 2011; McIntosh & Lobaugh, 2004) following a previously employed strategy (Gagnepain et al., 2017). We restricted our analysis to an independent domain-general inhibitory control mask derived from a meta-analytic conjunction analysis of 40 Stop-signal and 16 Think/No-Think fMRI studies (described below). Within this mask, we identified voxels where the BOLD signal from the main effect of Inhibit > Respond contrast depicted the largest joint covariance with the SSRT and SIF scores.

Specifically, Inhibit > Respond contrast values from each voxel of an MNI-normalised brain volume were aligned and stacked across participants into a brain activation matrix X, and SSRT and SIF scores were entered into a matrix Y. In both matrices, rows represented participants. We then individually mean-centred the X and Y matrices and normalised each row in the matrix X (representing each participant’s voxel activations) so that the row sum of squares equalled to one. Setting an equal variance of voxel activities across subjects ensured that the observed differences between participants were not due to overall differences in activation. Hereafter, a correlation of X and Y matrices produced a matrix R encoding the relationship between each voxel activity and behavioural scores across participants. We then applied a singular-value decomposition to the correlation matrix R to identify LVs that maximise the covariance between voxel activation (X) and behavioural measurements (Y). Each LV contained a single value for each participant representing the variance explained by the LV, and brain saliences, which are a weighted pattern across brain voxels representing the strength of the relationship between the BOLD signal and the behavioural scores.

To assess the statistical significance of each LV and the robustness of voxel saliences, we used 5000 permutation tests and 5000 bootstrapped resamples, respectively. By dividing each voxel’s initial salience by the standard error of its bootstrapped distribution, we obtained a bootstrapped standard ratio, equivalent to a z-score, to assess the significance of a given voxel. We thresholded the acquired scores at 1.96, corresponding to p < 0.05, two-tailed. The multivariate PLS analysis method does not require correction for multiple comparisons as it quantifies the relationship between the BOLD signal and behavioural scores in a single analytic step (McIntosh & Lobaugh, 2004).

##### Dynamic causal modelling (DCM) analysis

We conducted a DCM analysis (Friston et al., 2003) to determine the most likely inhibition-related interactions between domain-general inhibitory control areas in the right prefrontal cortex and domain-specific target areas. For the domain-specific target areas, we selected the left primary motor cortex (M1) and right hippocampus, based on our previous findings showing that stopping actions and stopping memories preferentially downregulates M1 and hippocampus, respectively (Schmitz et al., 2017).

DCM enables one to investigate hypothesised interactions among pre-defined brain regions by estimating the effective connectivity according to (1) the activity of other regions via intrinsic connections; (2) modulatory influences on connections arising through experimental manipulations; and (3) experimentally defined driving inputs to one or more of the regions (Friston et al., 2003). The intrinsic, modulatory, and driving inputs one specifies constitute the model structure assumed to represent the hypothesised neuronal network underlying the cognitive function of interest.

With DCM, a set of models can be defined that embody alternate hypotheses about the average connectivity and conditional moderation of connectivity. These models are inverted to the data and then compared in terms of the relative model evidence using Bayesian model selection (BMS). The differential model evidence from BMS indicates the probability that a given model is more likely to have generated the data than the other models and allows to infer both the presence and direction of modulatory connections. This can be estimated for individual models, or families of models that share critical features.

For the DCM analysis, we defined four regions of interest (ROIs): the right dorsolateral prefrontal cortex (rDLPFC), the right ventrolateral prefrontal cortex (rVLPFC), the right hippocampus, and the left M1. We obtained the rDLPFC and rVLPFC ROIs, centred at MNI coordinates 35, 45, 24 and 44, 21, −1, respectively, from an independent meta-analytic conjunction analysis (described below). We defined the M1 ROI (centred at MNI coordinates −33, −22, 46) from a group analysis (N = 30) of an independent M1 localiser study on different participants (Button Press > View contrast). We mapped the rDLPFC, rVLPFC, and M1 ROIs from the MNI space to participants’ native space. We manually traced the hippocampal ROIs in native space for each participant, using ITK-SNAP (www.itksnap.org; Yushkevich et al., 2006) and following established anatomical guidelines (Duvernoy et al., 2013; Pruessner et al., 2000). Within each subject-specific ROI, we identified all significant voxels (thresholded at p < 0.05, uncorrected for multiple comparisons) for that participant based on the main effect of interest, which included Stop, Go, No-Think, and Think conditions. Only the identified significant voxels were included in the final ROIs for the DCM analysis.

We performed the DCM analysis on participants’ native-space, unsmoothed brain images, to maximise the anatomical specificity of the hand-traced hippocampal ROI. We estimated a first-level GLM for each participant in their native space. The GLM model was closely similar to the first-level model defined for the univariate whole-brain analysis (see above). But in this new model, we concatenated all functional runs into a single run to form a single time series per participant. Because we concatenated the runs, we did not model conditions that started less than 24 s before the end of each run (apart from the very last run), and we did not use the SPM high-pass filtering and temporal autocorrelation options, but as additional regressors of no interest we included sines and cosines of up to three cycles per run to capture low-frequency drifts, and regressors modelling each run.

From each of the four ROIs, we extracted the first eigen-variate of the BOLD signal time-course, adjusted for effects of interest. Based on these data, we estimated and compared a null model with no modulatory connections and 72 models with modulatory connections (73 models in total) to test alternative hypotheses about how suppressing actions and memories modulate connectivity between the four ROIs (see Figure 6a). All 72 models with modulatory connections were variants of the same basic model with intrinsic bidirectional connections between all regions except no intrinsic connections between M1 and hippocampus, and with driving inputs from the Stop-signal (Stop and Go trials) and Think/No-Think (No-Think and Think trials) tasks into both rDLPFC and rVLPFC regions. Across models, we varied the modulatory influences on the intrinsic connections arising through Stop or No-Think trials.

We grouped the 72 models into three families differing according to whether the source-target modulation was bidirectional, top-down, or bottom-up. Within each family, we defined four subfamilies that differed according to how Stop and No-Think trials modulate the prefrontal control and inhibitory target pathways: independent modulation of target regions by rDLPFC and rVLPFC (testing the idea that two parallel inhibition pathways might exist); rDLPFC only modulation (testing the idea that only rDLPFC supports inhibition); rVLPFC only modulation (testing the idea that only rVLPFC supports inhibition); or modulation of both rDLPFC and rVLPFC (testing the idea that both contribute to inhibition). Within the four subfamilies, we defined further four subfamilies according to how Stop and No-Think trials modulate interactions between the rDLPFC and rVLPFC regions: no interactions; rVLPFC modulates rDLPFC; rDLPFC modulates rVLPFC; or bidirectional interaction between rDLPFC and rVLPFC.

Furthermore, within each subfamily, we defined two additional subfamilies according to whether Stop and No-Think trials modulate the prefrontal connectivity with the preferred targets (M1 when stopping actions and hippocampus when stopping memories) or with the non-preferred targets (hippocampus when stopping actions and M1 when stopping memories), testing the idea that inhibitory modulation must affect a task appropriate structure to model the data well.

We compared the model evidence for the 73 models (the null model and 72 models with modulatory connections) and the groups and subgroups of families across the 24 subjects using random-effects BMS (Penny et al., 2010; Stephan et al., 2010). BMS reports the exceedance probability, which is a probability that a given model, or family of models, is more likely than any other model or family tested, given the group data.

##### Multi-voxel pattern analysis

We performed multi-voxel pattern analysis (MVPA) to test whether action and memory inhibition share a common voxel activation pattern within an ROI. We used linear discriminant analysis (LDA) to classify voxel activity patterns within the same four ROIs that we used for the DCM analysis (rDLPFC, rVLPFC, right hippocampus, and left M1).

For each participant on their native-space unsmoothed brain images, we estimated a first-level GLM which was identical to the first-level model defined for the univariate whole-brain analysis (see above). The estimated beta weights of the voxels in each ROI were extracted and pre-whitened to construct noise normalized activity patterns for each event of interest (No-Think, Think, Stop, Go) within each of the eight functional fMRI runs.

To increase the reliability of pattern classification accuracy, we used a random subset approach (Diedrichsen et al., 2013). Specifically, for each ROI separately, we created up to 2000 unique subsets of randomly drawn 90% of ROI voxels (for smaller ROIs, there were less than 2000 possible combinations). We then applied the LDA on each subset and averaged the subset results to obtain the final classification accuracy for each ROI. We performed two types of pattern classification to identify domain-general and domain-specific components within each ROI.

For the domain-general component, we performed a cross-task classification. We trained the LDA classifier to distinguish Inhibit from Respond conditions in one modality (e.g. No-Think from Think) and tested whether the trained classifier could distinguish Inhibit from Respond in the other modality (e.g. Stop from Go). Both training and testing data consisted of two (conditions) by eight (runs) activation estimates for a set of voxels (e.g. 13 x 16 matrix for a set of 13 voxels). For training and testing sets separately, for each voxel, we z-scored the activity patterns across the 16 activation estimates setting the mean activity within each voxel to zero. This way, each voxel represented only the relative contribution of Inhibit vs Respond conditions within the Think/No-Think and Stop-signal tasks. For each ROI subset, we performed the LDA twice. The first classifier trained to discriminate Think from No-Think and returned the accuracy of distinguishing Stop from Go; the second classifier trained to discriminate Stop from Go and returned the accuracy of distinguishing Think from No-Think. The final score was the average classification accuracy of all subsets and the two classification variants (up to 2000 x 2) per ROI and subject.

For the domain-specific component, we trained and tested the LDA classifier to distinguish No-Think from Stop conditions. The input data consisted of two (conditions) by eight (runs) activation estimates for a set of voxels. We z-scored the activity patterns across voxels for each event of interest. Thus, the mean ROI activity for each event was zero, and each voxel represented only its relative contribution to the given event. That way, we accounted for the univariate intensity differences between No-Think and Stop conditions. For each ROI subset, we performed leave-one-run out cross-validated LDA and averaged the classification accuracies across the eight cross-validation folds. The final score was the average classification accuracy of all subsets and cross-validation folds (up to 2000 x 8) per ROI and subject.

At the group level, for each ROI, we performed one-tailed t-tests to assess the statistical significance of classification accuracy being above the 50% chance level. All tests were Bonferroni corrected for the number of ROIs.

##### A meta-analytic conjunction analysis of Stop-signal and Think/No-Think studies

To acquire an independent mask of brain areas involved in domain-general inhibitory control, we updated a previously published meta-analysis of Stop-signal and Think/No-Think fMRI studies (Guo et al., 2018). The study selection process and included studies are reported in detail in (Guo et al., 2018). From the original meta-analysis, we excluded the current dataset (Schmitz et al., 2017) and included a different within-subjects (but with each task performed on different days) Stop-signal and Think/No-Think study from our lab (Guo, 2017). Consequently, our analysis included 40 Stop-signal and 16 Think/No-Think studies. We focused the meta-analysis on the conjunction of Stop > Go No-Think > Think contrasts which we conducted using Activation Likelihood Estimation (ALE) with GingerALE v3.0.2 (http://www.brainmap.org/ale/; Eickhoff et al., 2012; Eickhoff et al., 2017; Eickhoff et al., 2009; Turkeltaub et al., 2012). We used the same settings as reported before (Guo et al., 2018). Specifically, we used a less conservative mask size, a non-additive ALE method, no additional FWHM, and cluster analysis peaks at all extrema. In addition, we set the coordinate space to MNI152.

First, we conducted separate meta-analyses of Stop > Go, No-Think > Think, and their pooled data using cluster-level FWE corrected inference (p < 0.05, cluster-forming threshold uncorrected p < 0.001, threshold permutations = 1000). We then submitted the obtained thresholded ALE maps from the three individual meta-analyses to a meta-analytic contrast analysis (Eickhoff et al., 2011), which produced the conjunction of the Stop > Go & No-Think > Think contrasts. We thresholded the conjunction results at voxel-wise uncorrected p < 0.001, with the p-value permutations of 10,000 iterations, and the minimum cluster volume of 200 mm^3^.

