## Supplemental Tables & Figures for "Dynamic targeting enables domain-general inhibitory control over action and thought by the prefrontal cortex"

### Supplementary Tables & Figures

#### Domain-general region overlap with the Cole-Anticevic brain-wide networks

To characterise these activations in relation to large-scale brain networks, we used a publicly available Cole-Anticevic brain-wide network partition (CAB-NP) (Ji et al., 2019). The CAB-NP was derived from resting-state fMRI data across the whole brain and used the Louvain community detection algorithm to assign parcellated cortical regions (Glasser et al., 2016) into 12 functional networks. We used the Connectome Workbench software (Marcus et al., 2011) to overlay our activations over the CAB-NP to estimate the parcel and network locations of our activation clusters.

For each estimated parcel we also identified its global variability coefficient (GVC), between-network variability coefficient (BVC) and network partition deviation numbers from Cocuzza et al. (2020) discovery and replication data sets. In Tables S1-S3 we present the identified parces for each of our activation clusters (Table 1 in the main text), their network assignment and center X, Y, Z coordinates, and GVC, BVC and deviation values averaged across Cocuzza et al. (2020) discovery and replication data sets. The top panels of Figures S1-S3 show our results clusters and estimated network assigments. Number of dots indicate to how many parcels within this network our cluster was assigned to. Dots were placed manually and are for the visualisation purposes only. The bottom panels show CAB-NP, our result maps in white and foci of our clusters. The foci were placed based on the MNI coordinates from the Table 1 of the main text.

##### Within-subjects Stop > Go & No-Think > Think conjunction clusters

**Table S1**

| Parcel | Network | X | Y | Z | GVC | BVC | deviation | Parcel description |
|---|---|---|---|---|---|---|---|---|
| **1. R VLPFC & Insula** | | | | | | | | |
| 254 | Frontoparietal | 53 | 19 | 13 | 0.3469 | 0.3486 | 0.6563 | R_Area_44 |
| 258 | Cingular-Opercular | 53 | 11 | 13 | 0.3534 | 0.3553 | 0.0000 | R_Rostral_Area_6 |
| 288 | Cingular-Opercular | 39 | 16 | 6 | 0.3717 | 0.3721 | 0.8907 | R_Frontal_Opercular_Area_4 |
| 289 | Cingular-Opercular | 38 | 13 | 1 | 0.3663 | 0.3691 | 0.1172 | R_Middle_Insular_Area |
| 291 | Frontoparietal | 33 | 26 | -4 | 0.3289 | 0.3299 | 0.3204 | R_Anterior_Ventral_Insular_Area |
| 349 | Cingular-Opercular | 38 | 28 | 4 | 0.3714 | 0.3724 | 0.0078 | R_Area_Frontal_Opercular |
| **Mean** | | | | | 0.3564 | 0.3579 | 0.3321 | |
| Cingular-Opercular | | | | | 0.3657 | 0.3672 | 0.2539 | |
| Frontoparietal | | | | | 0.3379 | 0.3393 | 0.4884 | |
| **2. Right IPL** | | | | | | | | |
| 205 | Cingular-Opercular | 63 | -37 | 27 | 0.3789 | 0.3818 | 0.1797 | R_PeriSylvian_Language_Area |
| 208 | Posterior Multimodal | 57 | -45 | 22 | 0.3609 | 0.3620 | 0.3282 | R_Superior_Temporal_Visual_Area |
| 328 | Cingular-Opercular | 60 | -30 | 38 | 0.3443 | 0.3446 | 0.0000 | R_Area_PF_Complex |
| 329 | Frontoparietal | 51 | -50 | 45 | 0.3351 | 0.3361 | 0.2501 | R_Area_PFm_Complex |
| **Mean** | | | | | 0.3548 | 0.3561 | 0.1895 | |
| **3. Right SMA** | | | | | | | | |
| 206 | Language | 8 | 19 | 64 | 0.3794 | 0.3819 | 0.1954 | R_Superior_Frontal_Language_Area |
| 224 | Cingular-Opercular | 20 | 7 | 66 | 0.3463 | 0.3477 | 0.5079 | R_Area_6m_anterior |
| 278 | Frontoparietal | 20 | 25 | 57 | 0.3812 | 0.3831 | 1.0000 | R_Superior_6-8_Transitional_Area |
| **Mean** | | | | | 0.3690 | 0.3709 | 0.5678 | |
| **4. Right DLPFC** | | | | | | | | |
| 264 | Cingular-Opercular | 36 | 41 | 30 | 0.3608 | 0.3625 | 0.0000 | R_Area_46 |
| 266 | Cingular-Opercular | 29 | 50 | 22 | 0.3750 | 0.3770 | 0.0547 | R_Area_9-46d |
| **Mean** | | | | | 0.3679 | 0.3698 | 0.0274 | |
| **5. Right Precentral** | | | | | | | | |
| 190 | Cingular-Opercular | 44 | -2 | 51 | 0.3653 | 0.3667 | 0.0391 | R_Frontal_Eye_Fields |
| 191 | Cingular-Opercular | 47 | 3 | 37 | 0.3618 | 0.3625 | 0.0782 | R_Premotor_Eye_Fields |
| 192 | Language | 49 | 2 | 47 | 0.3996 | 0.4007 | 0.0078 | R_Area_55b |
| 253 | Frontoparietal | 40 | 18 | 36 | 0.3347 | 0.3357 | 0.0782 | R_Area_IFJp |

| | | | | | | | | |
|---|---|---|---|---|---|---|---|---|
| **Mean** | | | | | 0.3654 | 0.3664 | 0.0508 | |
| **6. Left IPL** | | | | | | | | |
| 148 | Cingular-Opercular | -61 | -36 | 36 | 0.3635 | 0.3653 | 0.6875 | L_Area_PF_Complex |
| 149 | Frontoparietal | -50 | -56 | 44 | 0.3767 | 0.3799 | 0.1094 | L_Area_PFm_Complex |
| **Mean** | | | | | 0.3701 | 0.3726 | 0.3985 | |

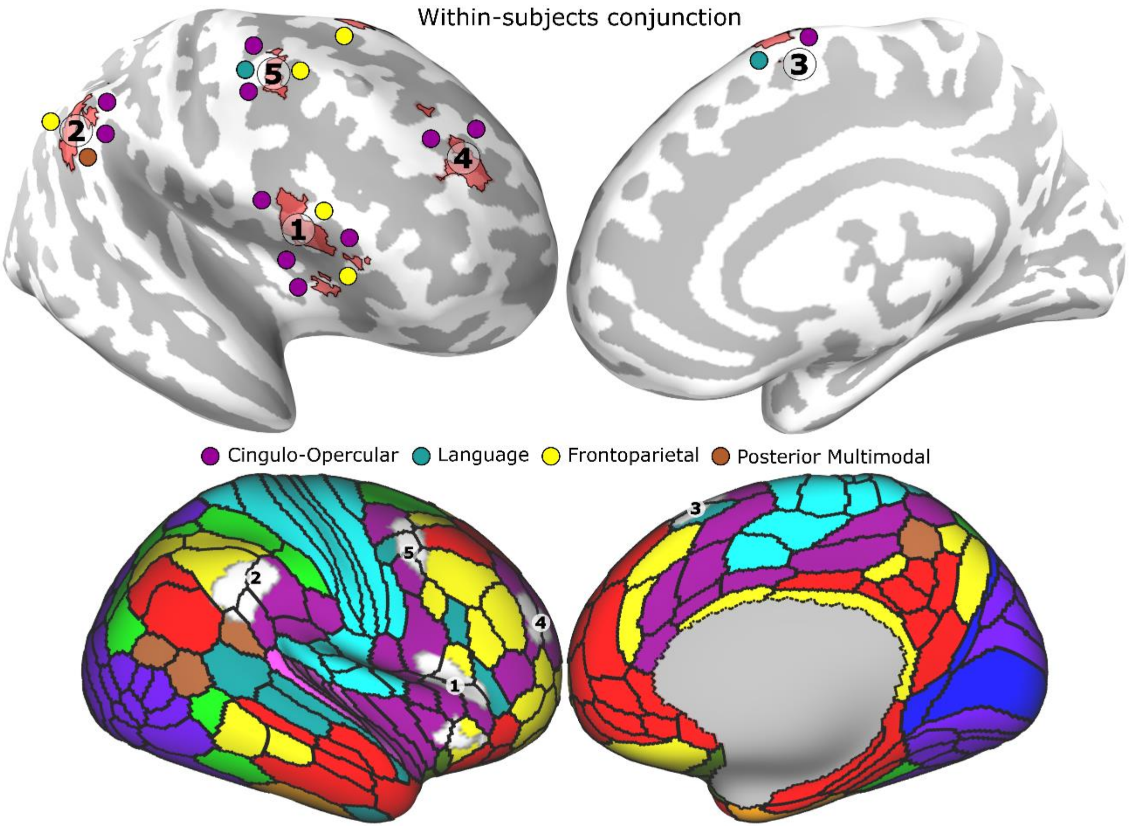


**Figure S1**

#### Meta-analysis Stop > Go & No-Think > Think conjunction clusters

**Table S2**

| Parcel | Network | X | Y | Z | GVC | BVC | deviation | Parcel description |
|---|---|---|---|---|---|---|---|---|
| **1. Right VLPFC & Insula** | | | | | | | | |
| 254 | Frontoparietal | 53 | 19 | 13 | 0.3469 | 0.3486 | 0.6563 | R_Area_44 |
| 288 | Cingular-Opercular | 39 | 16 | 6 | 0.3717 | 0.3721 | 0.8907 | R_Frontal_Opercular_Area_4 |
| 289 | Cingular-Opercular | 38 | 13 | 1 | 0.3663 | 0.3691 | 0.1172 | R_Middle_Insular_Area |
| 291 | Frontoparietal | 33 | 26 | -4 | 0.3289 | 0.3299 | 0.3204 | R_Anterior_Ventral_Insular_Area |
| 349 | Cingular-Opercular | 38 | 28 | 4 | 0.3714 | 0.3724 | 0.0078 | R_Area_Frontal_Opercular |
| **Mean** | | | | | 0.3570 | 0.3584 | 0.3985 | |
| Cingular-Opercular | | | | | 0.3698 | 0.3712 | 0.3386 | |
| Frontoparietal | | | | | 0.3379 | 0.3393 | 0.4884 | |
| **2. Right/Left SMA** | | | | | | | | |
| 206 | Language | 8 | 19 | 64 | 0.3794 | 0.3819 | 0.1954 | R_Superior_Frontal_Language_Area |
| 223 | Cingular-Opercular | 6 | 6 | 58 | 0.3928 | 0.3951 | 0.0625 | R_Supplementary_And_Cingulate_Eye |
| 224 | Cingular-Opercular | 20 | 7 | 66 | 0.3463 | 0.3477 | 0.5079 | R_Area_6m_anterior |
| 243 | Frontoparietal | 5 | 32 | 46 | 0.3692 | 0.3698 | 0.2344 | R_Area_8BM |
| 278 | Frontoparietal | 20 | 25 | 57 | 0.3812 | 0.3831 | 1.0000 | R_Superior_6-8_Transitional_Area |
| **Mean** | | | | | 0.3738 | 0.3755 | 0.4000 | |
| **3. Left VLPFC & Insula** | | | | | | | | |
| 109 | Cingular-Opercular | -37 | 11 | 3 | 0.3519 | 0.3552 | 0.2422 | L_Middle_Insular_Area |
| 111 | Frontoparietal | -31 | 25 | -3 | 0.3805 | 0.3847 | 0.3438 | L_Anterior_Ventral_Insular_Area |
| **Mean** | | | | | 0.3662 | 0.3700 | 0.2930 | |
| **4. Right IPL** | | | | | | | | |
| 205 | Cingular-Opercular | 63 | -37 | 27 | 0.3789 | 0.3818 | 0.1797 | R_PeriSylvian_Language_Area |
| 208 | Posterior Multimodal | 57 | -45 | 22 | 0.3609 | 0.3620 | 0.3282 | R_Superior_Temporal_Visual_Area |
| 328 | Cingular-Opercular | 60 | -30 | 38 | 0.3443 | 0.3446 | 0.0000 | R_Area_PF_Complex |
| 329 | Frontoparietal | 51 | -50 | 45 | 0.3351 | 0.3361 | 0.2501 | R_Area_PFm_Complex |

| | | | | | | | | |
|---|---|---|---|---|---|---|---|---|
| **Mean** | | | | | 0.3548 | 0.3561 | 0.1895 | |
| **5. Right ACC** | | | | | | | | |
| 239 | Cingular-Opercular | 4 | 19 | 32 | 0.3716 | 0.3705 | 0.8047 | R_Anterior_24_prime |
| 240 | Cingular-Opercular | 9 | 15 | 39 | 0.3795 | 0.3783 | 0.5782 | R_Area_p32_prime |
| 243 | Frontoparietal | 5 | 32 | 46 | 0.3692 | 0.3698 | 0.2344 | R_Area_8BM |
| 359 | Cingular-Opercular | 9 | 29 | 30 | 0.3592 | 0.3595 | 0.9922 | R_Area_anterior_32_prime |
| **Mean** | | | | | 0.3699 | 0.3695 | 0.6524 | |
| **6. Right DLPFC** | | | | | | | | |
| 264 | Cingular-Opercular | 36 | 41 | 30 | 0.3608 | 0.3625 | 0.0000 | R_Area_46 |
| 266 | Cingular-Opercular | 29 | 50 | 22 | 0.3750 | 0.3770 | 0.0547 | R_Area_9-46d |
| **Mean** | | | | | 0.3679 | 0.3698 | 0.0274 | |
| **7. Basal ganglia** | | | | | | | | |
| **8. Left IPL** | | | | | | | | |
| 148 | Cingular-Opercular | -61 | -36 | 36 | 0.3635 | 0.3653 | 0.6875 | L_Area_PF_Complex |
| 149 | Frontoparietal | -50 | -56 | 44 | 0.3767 | 0.3799 | 0.1094 | L_Area_PFm_Complex |
| **Mean** | | | | | 0.3701 | 0.3726 | 0.3985 | |
| **9. Right Precentral** | | | | | | | | |
| 190 | Cingular-Opercular | 44 | -2 | 51 | 0.3653 | 0.3667 | 0.0391 | R_Frontal_Eye_Fields |
| 192 | Language | 49 | 2 | 47 | 0.3996 | 0.4007 | 0.0078 | R_Area_55b |
| **Mean** | | | | | 0.3825 | 0.3837 | 0.0235 | |
| **10. Right SPL** | | | | | | | | |
| 297 | Dorsal Attention | 38 | -38 | 44 | 0.3888 | 0.3899 | 0.9610 | R_Anterior_IntraParietal_Area |
| 324 | Frontoparietal | 42 | -42 | 46 | 0.3960 | 0.4011 | 0.2500 | R_Area_IntraParietal_2 |
| **Mean** | | | | | 0.3924 | 0.3955 | 0.6055 | |

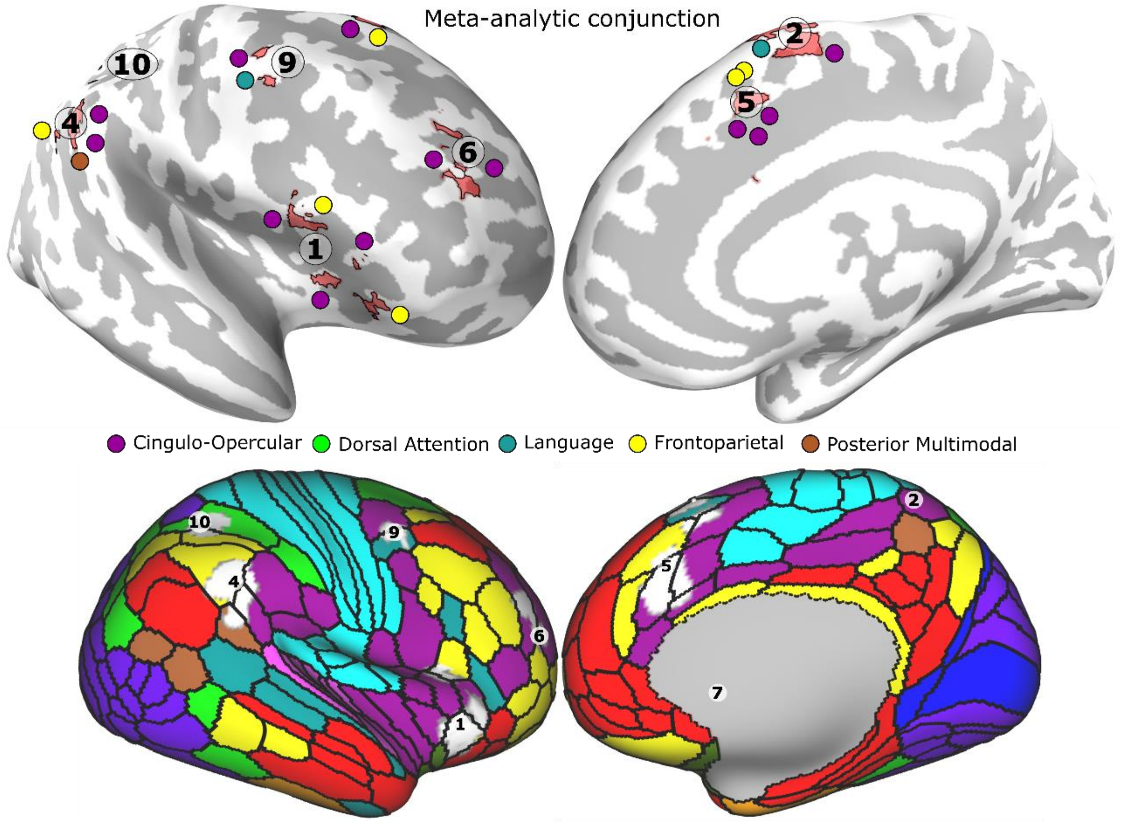


**Figure S2**

#### Conjunction of the within-subjects conjunction & meta-analysis conjuncion clusters

**Table S3**

| Parcel | Network | X | Y | Z | GVC | BVC | deviation | Parcel description |
|---|---|---|---|---|---|---|---|---|
| **1. Right VLPFC & Insula** | | | | | | | | |
| 254 | Frontoparietal | 53 | 19 | 13 | 0.3469 | 0.3486 | 0.6563 | R_Area_44 |
| 288 | Cingular-Opercular | 39 | 16 | 6 | 0.3717 | 0.3721 | 0.8907 | R_Frontal_Opercular_Area_4 |
| 289 | Cingular-Opercular | 38 | 13 | 1 | 0.3663 | 0.3691 | 0.1172 | R_Middle_Insular_Area |
| 291 | Frontoparietal | 33 | 26 | -4 | 0.3289 | 0.3299 | 0.3204 | R_Anterior_Ventral_Insular_Area |

| | | | | | | | | |
|---|---|---|---|---|---|---|---|---|
| 349 | Cingular-Opercular | 38 | 28 | 4 | 0.3714 | 0.3724 | 0.0078 | R_Area_Frontal_Opercular |
| **Mean** | | | | | 0.3570 | 0.3584 | 0.3985 | |
| Cingular-Opercular | | | | | 0.3698 | 0.3712 | 0.3386 | |
| Frontoparietal | | | | | 0.3379 | 0.3393 | 0.4884 | |
| **2. Right IPL** | | | | | | | | |
| 205 | Cingular-Opercular | 63 | -37 | 27 | 0.3789 | 0.3818 | 0.1797 | R_PeriSylvian_Language_Area |
| 208 | Posterior Multimodal | 57 | -45 | 22 | 0.3609 | 0.3620 | 0.3282 | R_Superior_Temporal_Visual_Area |
| 328 | Cingular-Opercular | 60 | -30 | 38 | 0.3443 | 0.3446 | 0.0000 | R_Area_PF_Complex |
| 329 | Frontoparietal | 51 | -50 | 45 | 0.3351 | 0.3361 | 0.2501 | R_Area_PFm_Complex |
| **Mean** | | | | | 0.3548 | 0.3561 | 0.1895 | |
| **3. Right SMA** | | | | | | | | |
| 206 | Language | 8 | 19 | 64 | 0.3794 | 0.3819 | 0.1954 | R_Superior_Frontal_Language_Area |
| 278 | Frontoparietal | 20 | 25 | 57 | 0.3812 | 0.3831 | 1.0000 | R_Superior_6-8_Transitional_Area |
| 224 | Cingular-Opercular | 20 | 7 | 66 | 0.3463 | 0.3477 | 0.5079 | R_Area_6m_anterior |
| **Mean** | | | | | 0.3690 | 0.3709 | 0.5678 | |
| **4. Right DLPFC** | | | | | | | | |
| 264 | Cingular-Opercular | 36 | 41 | 30 | 0.3608 | 0.3625 | 0.0000 | R_Area_46 |
| 266 | Cingular-Opercular | 29 | 50 | 22 | 0.3750 | 0.3770 | 0.0547 | R_Area_9-46d |
| **Mean** | | | | | 0.3679 | 0.3698 | 0.0274 | |
| **5. Left IPL** | | | | | | | | |
| 148 | Cingular-Opercular | -61 | -36 | 36 | 0.3635 | 0.3653 | 0.6875 | L_Area_PF_Complex |
| 149 | Frontoparietal | -50 | -56 | 44 | 0.3767 | 0.3799 | 0.1094 | L_Area_PFm_Complex |
| **Mean** | | | | | 0.3701 | 0.3726 | 0.3985 | |
| **6. Right Precentral** | | | | | | | | |
| 190 | Cingular-Opercular | 44 | -2 | 51 | 0.3653 | 0.3667 | 0.0391 | R_Frontal_Eye_Fields |
| 192 | Language | 49 | 2 | 47 | 0.3996 | 0.4007 | 0.0078 | R_Area_55b |
| **Mean** | | | | | 0.3825 | 0.3837 | 0.0235 | |

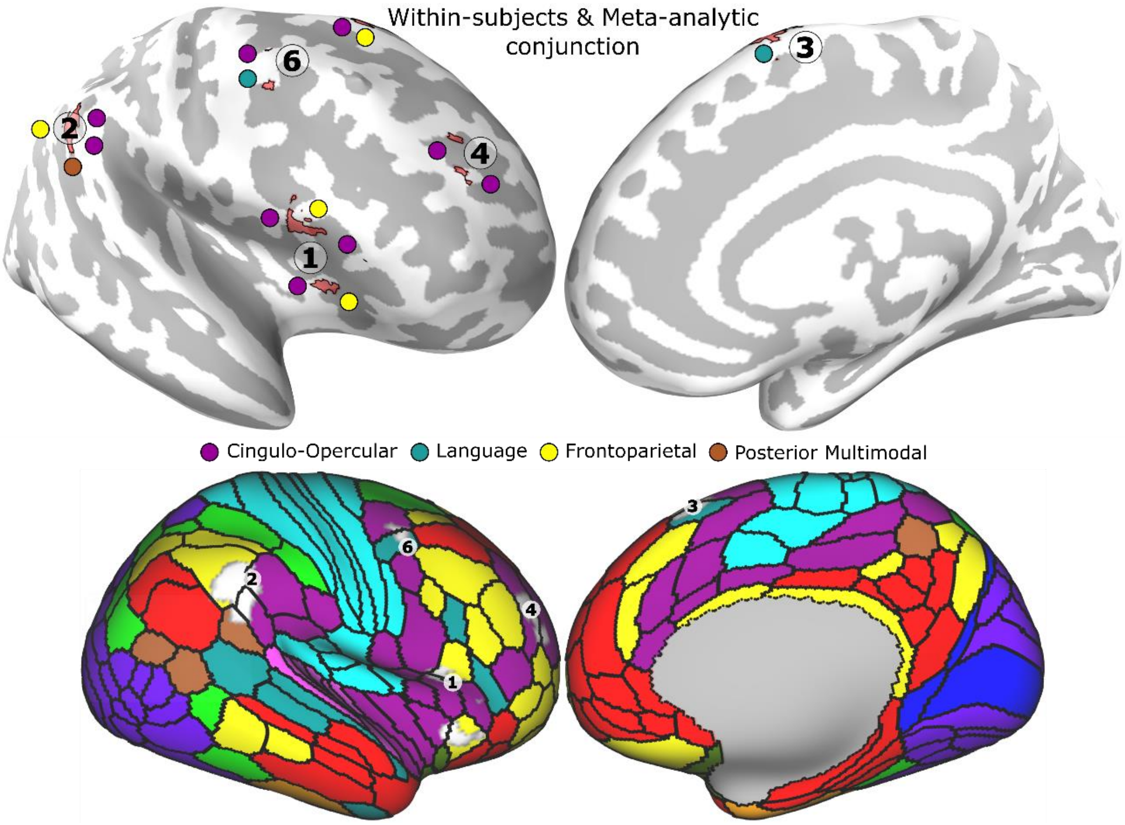


**Figure S3**

#### Conflict reduction

Figures S4A and S5A show how accurately a classifier trained to distinguish Stop from Go conditions, classifies No-Think as Stop condition for each of the 8 fMRI task runs in the rDLPFC and rVLPFC, respectively.

Figures S4B and S5B show correlation between the suppression induced forgetting (SIF) and slope of the classification accuracy over runs (measured as the accuracy slope divided by the accuracy of the first run), in the rDLPFC and rVLPFC, respectively.

Figures S4C and S5C show correlation between the stop-signal reaction time (SSRT) and slope of the classification accuracy over runs, in the rDLPFC and rVLPFC, respectively.

##### rDLPFC

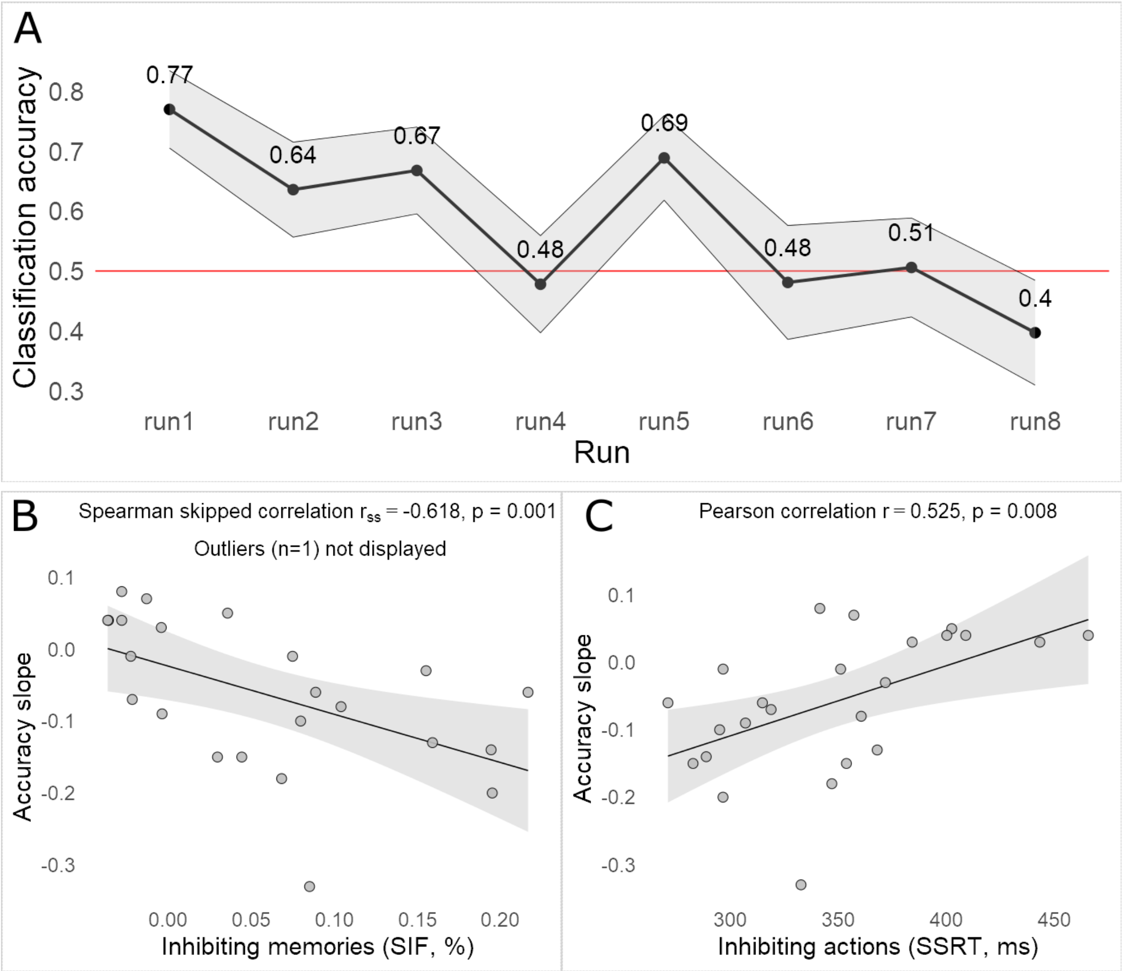


**Figure S4**

rVLPFC

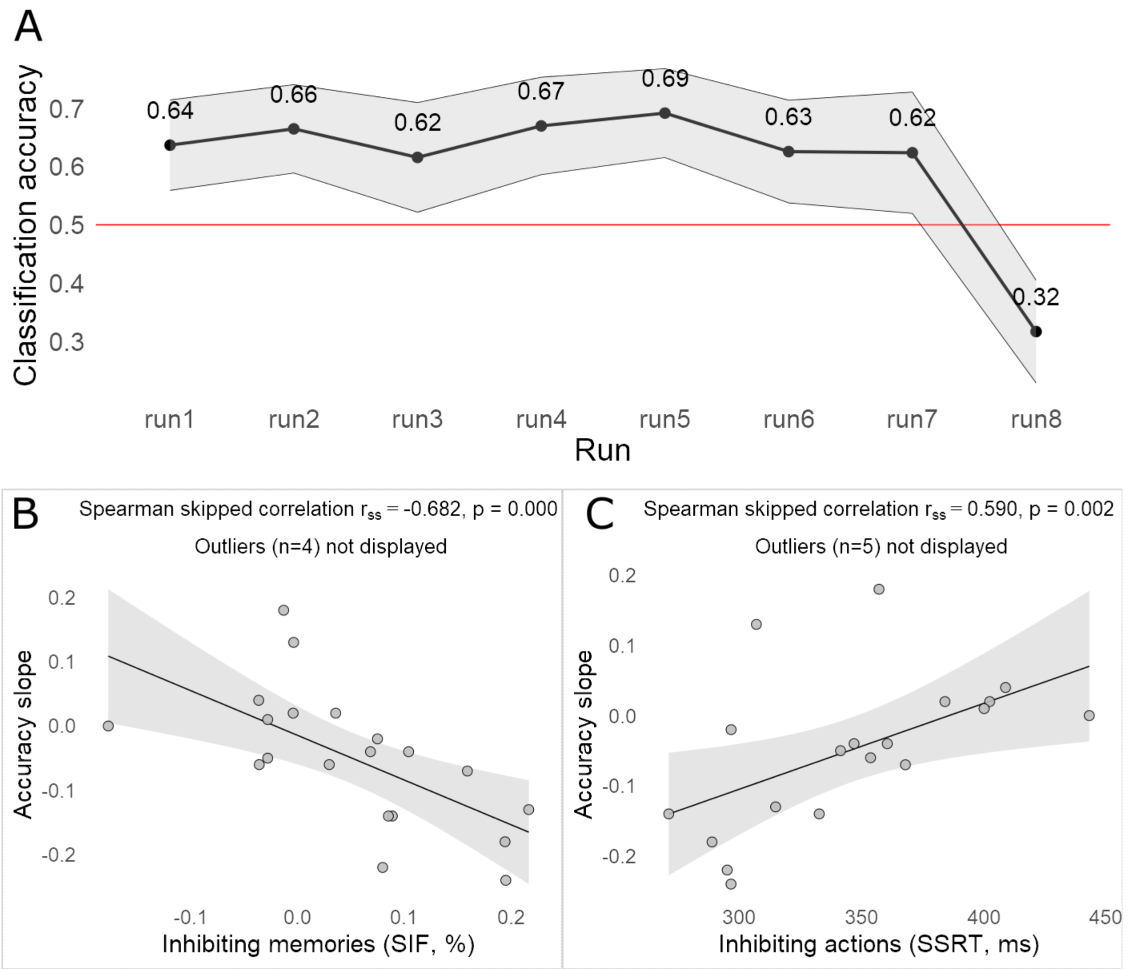


**Figure S5.**
